# Ribonuclease Inhibitor and Angiogenin collaboratively regulate cell-type-specific global translation

**DOI:** 10.1101/2024.03.29.586999

**Authors:** Martina Stillinovic, Mayuresh Anant Sarangdhar, Nicola Andina, Aubry Tardivel, Frédéric Greub, Giuseppe Bombaci, Camille Ansermet, Manfred Heller, Adrian Keogh, Irene Keller, Anne Angelillo-Scherrer, Ramanjaneyulu Allam

## Abstract

Translation of mRNAs is a fundamental process that occurs in all cell-types of multicellular organisms. Conventionally, it has been considered a default step in gene expression, lacking specific regulation. However, recent studies have documented that certain mRNAs exhibit cell-type-specific translation^1–3^. Despite this, it remains unclear whether global translation is controlled in a cell-type-specific manner. Here we report that a ribosome-associated protein ribonuclease inhibitor-1 (RNH1) and its binding partner Angiogenin (ANG) collaboratively regulates cell-type-specific global translation. By employing human cell-lines and mouse models, we found that deletion of RNH1 decreases global translation selectively in hematopoietic origin cells but not in the non-hematopoietic origin cells. RNH1 mediated such cell-type-specific translation is mechanistically linked to ANG. We found that ANG, which is known to regulate ribosomal biogenesis^4^, is predominantly expressed in non-hematopoietic origin cells and absent in hematopoietic origin cells. ANG safeguards the non-hematopoietic origin cells from RNH1-knockout-mediated translation defects by upregulating ribosomal biogenesis. Further, we discovered that RNH1 controls the translation of ribosomal protein (RP) transcripts and influences mRNA circularization. Collectively, this study unravels the existence of cell-type-specific global translation regulators and highlights the complex translation regulation in vertebrates.

## Main Text

Gene expression diversifies the phenotype and functions of cells in multicellular organisms. Primarily, gene expression is regulated at the transcriptional level through the action of transcription factors ^5^. While translation is crucial for decoding information from mRNAs to produce proteins, it has traditionally been seen as a default process without direct regulation of specific gene expression. However, recent studies indicate that translational control plays a crucial role in fine-tuning protein synthesis and modulating the expression of specific mRNAs ^1^. For example, a mutation in the ribosomal protein L38 (*RPL38*) gene in mice deregulates the translation of a subset of homeobox mRNAs, leading to tissue-specific patterning defects ^2^. Further, mutations in several ribosomal protein (RP) genes, such as *RPS19, RPL5, RPL11, RPL35A, RPS10, RPS17, RPS24,* cause Diamond Blackfan anemia (DBA)^6^. DBA is characterized by bone marrow failure, anemia and craniofacial malformations ^7^. Studies have shown that decreased translation of certain mRNAs involved in erythroid differentiation is responsible for the anemia phenotype in DBA ^3,8^. RPs are core components of ribosomes and are present in every cell. Mutations in their genes should cause global phenotype, but what accounts for these tissue-specific phenotypes after translational defects is not completely understood. Several mechanisms have been proposed to address this and are matter of debate ^9–12^. Although specific mRNA translation adds another layer of gene expression regulation, it is not yet known whether global translation is controlled in a cell-type-specific manner.

Ribonuclease inhibitor 1 (RI or RNH1) also known as Ribonuclease/angiogenin inhibitor is a ubiquitously expressed protein in mammalian cells ^13,14^. Phylogenetically, *RNH1* has no apparent homologues in invertebrates and lower vertebrates (**Extended Data Fig. 1**), suggesting it is a higher vertebrate specific gene. It evolved via exon duplication and is conserved among mammals^15^. RNH1 is known to bind and inhibits pancreatic-type ribonucleases such as RNase 1, RNase 2, RNase 4 and angiogenin (ANG, also known as RNase 5)^13^. RNH1 is composed entirely of leucine-rich repeats (LRRs) and displays vast surface areas that foster protein-protein and protein-ligand interactions ^16^. Consistent with this, RNH1 has been reported to have other biological functions besides its RNase inhibitor role, such as binding and inhibiting the ER stress sensor IRE1 activity ^17^, involvement in cancer growth and metastasis ^18^, tRNA degradation ^19^, microRNA (miR-21) processing ^20^, and inhibiting inflammasome activation ^16^. Previously, we have reported that RNH1 is a ribosome-associated protein that regulates GATA1 translation, and responsible for defects in embryonic erythropoiesis in RNH1 deficient mice ^21^. In line with this, recent human molecular genetics studies have further revealed that mutations in *RNH1* gene lead to severe phenotypes such as global developmental delay, embryonic death, inflammation, and macrocytic anemia ^22^. Collectively, these findings strongly suggest that RNH1 is a multifunctional molecule crucial for mammalian survival ^23^. In this study, we employed a combination of human cell-lines and mouse models, along with biochemical experiments, to establish the critical role of RNH1 in hematopoietic-specific translation. We found that RNH1 via its binding partner ANG, confers cell-type-specific global translation regulation. Additionally, we discovered that RNH1 controls the translation of RP mRNAs and influences mRNA circularization. These findings provide valuable insights into the intricate regulatory mechanisms governing cell-type-specific translation processes.

### The loss of RNH1 predominantly reduces translation in hematopoietic origin cells

RNH1 is a ribosome-associated protein and deletion of RNH1 in a human erythroleukemic cell line (K562 cells) has been shown to result in reduced polysomes levels ^21^ (**Extended Data Fig. 2A**). However, the detailed mechanism/s of RNH1 mediated translation regulation remains unknown. To investigate this, RNH1 was knocked out of HEK293T cells, and the global translation status was checked by polysome analysis (**Fig. 1a**). Surprisingly, polysome levels were not altered in RNH1 knockout (KO) cells, suggesting that RNH1 might regulate cell-type-specific translation. Next, RNH1 was knocked out in several randomly selected cell lines of different origins, and global translation status was checked by polysome analysis (**Fig. 1a, b**). Interestingly, cell lines from hematopoietic origin (THP1, MOLM13 and Jurkat) consistently showed a large reduction in polysome levels compared to their corresponding WT cells (**Fig. 1b**). However, polysome levels were not changed in RNH1 KO non-hematopoietic origin cells-lines such as HeLa, HaCaT, and SHSY5Y compared to corresponding WT cells (**Fig. 1a and Extended Data Fig. 2b**). To our knowledge, such cell-type-specific global translation defect has not been reported previously in mammalian cells. While mutations in RPs lead to cell-type-specific functional defects ^6^, their knockdown dysregulates global translation in various cell-types of different origin. For example, the *RPS19* gene is frequently mutated in DBA patients, and its knockdown leads to global translational defects in several cell-types including both hematopoietic and non-hematopoietic origin cells (HeLa, embryonic stem cells, different erythroid cell lines, and progenitor cells)^3,8,24–27^. These results suggest that, unlike RP gene mutations, loss of RNH1 primarily leads to translation defects in hematopoietic origin cells but not in non-hematopoietic origin cells.

**Fig 1.**
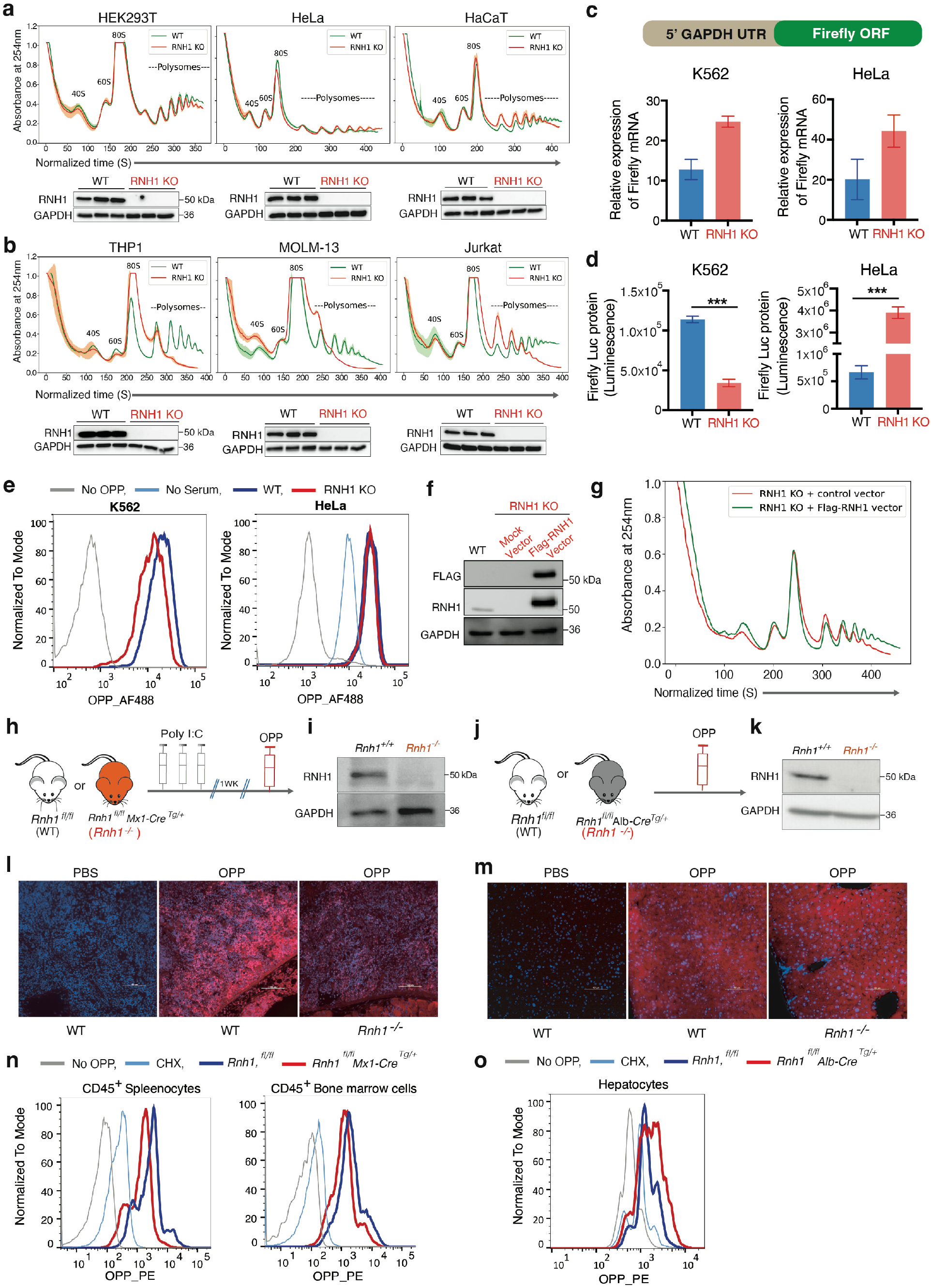
Loss of RNH1 decreases translation specifically in hematopoietic origin cells. (**a and b**) Sucrose gradient polysome profiles for wildtype (WT) and corresponding RNH1 knockout (RNH1 KO) non-hematopoietic (a) and hematopoietic (b) origin cells (N=3). Arrow shows the direction of the sucrose gradient from low to high density. Mean value of absorbance from three independent experiments plotted with the standard deviation (upper panel). Total protein lysates of WT and RNH1 KO cells were analyzed by western blot with the indicated antibodies. Blots are representative of three independent experiments (lower panel). (**c and d**) Schematics of luciferase expressing plasmid with GAPDH 5’UTR (upper panel). WT and RNH1 KO HeLa and K562 cells were transfected with luciferase expressing plasmid. Cells were analysed for firefly mRNAs by qRT-PCR, normalized to 18S rRNA expression (c) and luciferase protein expression by luciferase assay (d). Data are shown as mean ± SEM and are representative of 3 independent experiments. (**e**) WT or RNH1 KO K562 and HeLa cells were incubated for 1h with OPP and FACS analysis were performed to measure OPP incorporation. Representative histograms were shown for OPP fluorescence. Data is representative of 3 independent experiments. (**f and g**) RNH1 is ectopically expressed in RNH1 KO K562 cells. Cell lysates were analyzed by western blot with the indicated antibodies. Blots are representative of two independent experiments (f). Sucrose gradient polysome profiles for control and FLAG-RNH1 expressing RNH1 KO K562 cells. Data is representative of 3 independent experiments (g). (**h-m**) OPP incorporation assay was performed after 1h of intraperitoneal injection with PBS or OPP (50mg/kg) in WT (*Rnh1 ^fl/fl^*), hematopoietic-specific *Rnh1 deficient* (*Rnh1 ^fl/fl^, Mx-Cre^+^*) and liver-specific *Rnh1 deficient* (*Rnh1 ^fl/fl^, Alb-Cre^+^*) mice. Schematics showing Mx-Cre model, where *Rnh1* was excised by giving three rounds of 200μg polyIC using intraperitoneal injections to *Rnh1^fl/fl^* (WT) and *Rnh1^fl/fl^Mx1-Cre^+^* (*Rnh1^-/-^*) mice. After one week, mice were injected with OPP and organs were harvested after 1 hr (h). Spleen cell lysates of WT and *Rnh1^-/-^* cells were analyzed by western blot with the indicated antibodies. Blots are representative of three independent experiments (i). Schematics showing in *Rnh1^fl/fl^* (WT) and *Rnh1^fl/fl^ Alb-Cre^+^*(*Rnh1^-/-^*) mice were injected with OPP and organs were harvested after 1 hr (j). Liver cell lysates of WT and *Rnh1^-/-^* cells were analyzed by western blot with the indicated antibodies. Blots are representative of three independent experiments (k). OPP incorporated spleen (l) and liver (m) tissue sections were stained by CuAAC with TMR-azide and Hoechst (n =3 mice for each genotype) (scale 100µM). (**n and o**) WT or *Rnh1^-/-^* splenocytes, bone marrow cells (n) and hepatocytes (o) were incubated for 1h with OPP and FACS analysis was performed to measure OPP incorporation. Representative histograms were shown for OPP fluorescence (n =2 mice for each genotype). CHX (Cycloheximide).

To confirm above polysome results, luciferase reporter assay was performed to check the translation. HeLa WT and RNH1 KO cells, as well as K562 WT and RNH1 KO cells were transfected with a plasmid containing luciferase reporter gene having GAPDH 5’UTR (**Fig. 1c**). By maintaining same levels of luciferase mRNA in WT and RNH1 KO (**Fig. 1c**), we observed enhanced luciferase protein levels in HeLa RNH1 KO (**Fig. 1d**). On the contrary, K562 RNH1 KO cells showed a significant reduction in translation of luciferase reporter (**Fig. 1d**). Similar results were obtained with another luciferase reporter having beta actin (ACTB) 5’UTR (**Extended Data Fig. 2C**). Both ACTB and GAPDH 5’UTRs are simple UTRs without any secondary structure ^3^. Thus, this assay ruled out any contribution of the complexity of 5’UTRs in RNH1 mediated translation regulation. To further ascertain RNH1 mediated specific translation regulation, nascent polypeptide labeling via O-propargyl-puromycin (OPP) was performed. OPP is an analog of puromycin. It gets incorporated into newly translating proteins and can be detected by copper(I)-catalyzed azide-alkyne cycloaddition ^28^. Supporting the above results, flow cytometry analysis showed reduced incorporation of OPP in K562 RNH1 KO cells, but not in HeLa RNH1 KO cells, compared to their respective WT cells (**Fig. 1e**). Furthermore, ectopic expression of RNH1 in RNH1 KO K562 cells was able to rescue the polysome level defects observed in K562 RNH1 KO (**Fig. 1f and g**). Altogether, these results confirm that loss of RNH1 reduces global translation only in cells of hematopoietic origin.

Differences in cell size and ribosomal gene expression between hematopoietic and non-hematopoietic origin cell line might account for the observed translation defects ^29^. So, we checked cell size and ribosomal gene expression in these cell lines. These parameters varied across cell lines and none of them clearly separated hematopoietic cells from non-hematopoietic origin cell lines (**Extended Data Fig. 3a-c**). Thus, cell size and ribosomal gene expression could not account for the RNH1 mediated global translation defects in hematopoietic cell lines. It is known that RNH1 interacts with Angiogenin (ANG) and inhibits its functions. Under stress situations, ANG mediates translation repression by degradation of tRNAs to form tiRNAs in the cytoplasm ^19^. Therefore, we checked the possibility of increased tRNA degradation in RNH1 KO cells by assessing the levels of tiRNAs. Surprisingly, we did not find an increase in tiRNAs in the RNH1-KO cells at steady state without stress conditions (**Extended Data Fig. 3d**). However, under stressed conditions, RNH1 KO K562 cells showed a minor increase in tiRNA production compared to WT (**Extended Data Fig. 3d**). This corroborates with a previous study where overexpression of ANG in PC12 cells did not produce tiRNAs when cells were not stressed. However, under stress, ANG overexpressing cells synthesized larger amounts of tiRNAs, even with mild stress ^30^. Thus, under steady-state situations, we observed a perturbation of translation in RNH1 KO hematopoietic cells, along with unchanged tiRNA levels. These results exclude involvement of tRNA degradation in RNH1 mediated cell-type-specific translation regulation.

### RNH1 deficiency in mice specifically reduces the translation in hematopoietic cells

To confirm the RNH1-mediated hematopoietic-specific translation regulation *in vivo,* OPP was injected into WT (*Rnh1^fl/fl^*), hematopoietic *Rnh1* deficient (*Rnh1^fl/fl^, Mx-Cre^Tg/+^*) (**Fig. 1h** and **i**), and liver specific *Rnh1* deficient (*Rnh1^fl/fl^, Alb-Cre^Tg/+^*) mice (**Fig. 1j** and **k**). We refer *Rnh1* deficient mice *as Rnh1^-/-^*. Interestingly, corroborating our *in vitro* cell-line experiments, the spleen of *Rnh1^-/-^* mice showed reduced incorporation of OPP compared to WT (**Fig. 1l**), indicating translation reduction. On the contrary, the liver of *Rnh1^-/-^* mice did not show translation repression (**Fig. 1m**). Further, OPP incorporation assay was performed using hematopoietic cells isolated from bone marrow and spleen, and hepatocytes from liver. In agreements with *in vivo* results, translation was decreased in mouse hematopoietic cells (**Fig. 1n**), but not in hepatocytes (**Fig. 1o**). Thus, using multiple complementary approaches, our results demonstrated that RNH1 specifically regulates global translation in hematopoietic cells *in vivo* and *in vitro*. These results also suggest that RNH1-mediated hematopoietic translation is species-independent and occurs in both human and mice.

### RNH1 regulates translation of mRNAs encoding Ribosomal Proteins

To understand how RNH1 mediates hematopoietic-specific translation, we first examined its expression in different hematopoietic and non-hematopoietic cells lines. However, there was no major difference in RNH1 protein levels in both cell-types that could explain the above observed translational defects (**Fig. 2a**). Next, to gain insights into the molecular clues involved in RNH1-mediated cell-type-specific translation regulation, we performed polysome-associated RNA sequencing (polysome-seq) using polysomal RNA from K562 WT and RNH1 KO cells (**Fig. 2b**) and compared it with bulk RNA-sequencing (total-seq) data (**Supplementary Table. 1**) ^21^. Polysome-seq analysis revealed that mRNAs encoding RPs were significantly less associated with polysomes in RNH1 KO K562 cells compared to WT cells (**Fig. 2c**), indicating reduced translation of RPs mRNAs in RNH1 KO. On the contrary, total RNA-seq did not show any changes in the expression levels of RPs mRNAs between WT and RNH1 KO cells (**Fig. 2d**). This suggests that the translation of RPs mRNAs is perturbed in RNH1 KO hematopoietic cells without altering their mRNA levels. Next, translation activity (TA) was calculated as polysomal RNA/ total RNA. As expected, RPs mRNAs showed high expression and were more translationally active in WT cells (**Fig. 2e**). In RNH1 KO K562 cells, the TA of RPs mRNAs was reduced without changes in RNA expression levels (**Fig. 2f**), confirming the translation repression of RPs mRNAs in RNH1 KO cells. Additionally, we implemented analysis of translational activity (ANOTA), which applies per-gene analysis of partial variance with variance shrinkage ^31^. ANOTA identified a stringent set of 262 genes that were translationally deregulated in K562 RNH1 KO cells, including several RPs mRNAs (**Supplementary Table. 2**). This further confidently reiterates that the total RNA level of RPs mRNAs is not changed in WT and KO K562 cells, but their polysome-associated RNA levels decreased significantly in RNH1 KO K562 cells (**Extended Data Fig. 4a and b**). TA of genes retained after ANOTA analysis also showed that WT cells have high expression of RPs mRNAs and large TA, while in RNH1 KO, TA of RPs mRNAs decreased without change in their RNA levels (**Extended Data Fig. 4c and d**). We selected genes with high expression and large TA from WT (1549) and KO (884) (genes in the upper right quadrant of plots in **Fig. 2e and f**) and performed gene ontology (GO) analysis. The enrichment of gene ontology terms in WT and KO showed that terms like “translation”, “cytoplasmic translation” were significantly less represented in RNH1 KO compared to WT (**Fig. 2g**). This further suggests that RNH1 KO K562 cells have reduced translation activity. To confirm this, cell lysates of K562 and HeLa with WT and RNH1 KO genotypes were analyzed by western blots for representative RPs. As expected, K562 RNH1 KO cells showed reduced levels of RPS19 and RPS3 proteins compared to K562 WT cells. On the contrary, RPS19 and RPS3 levels in HeLa RNH1 KO cells did not decrease (**Fig. 2h**). Similarly, RPS19 and RPS3 levels were reduced in *Rnh1^-/-^* mice bone marrow cells but not in the liver compared to WT control mice (**Fig. 2i**). These results suggest RNH1 is involved in RP gene translation.

**Fig 2.**
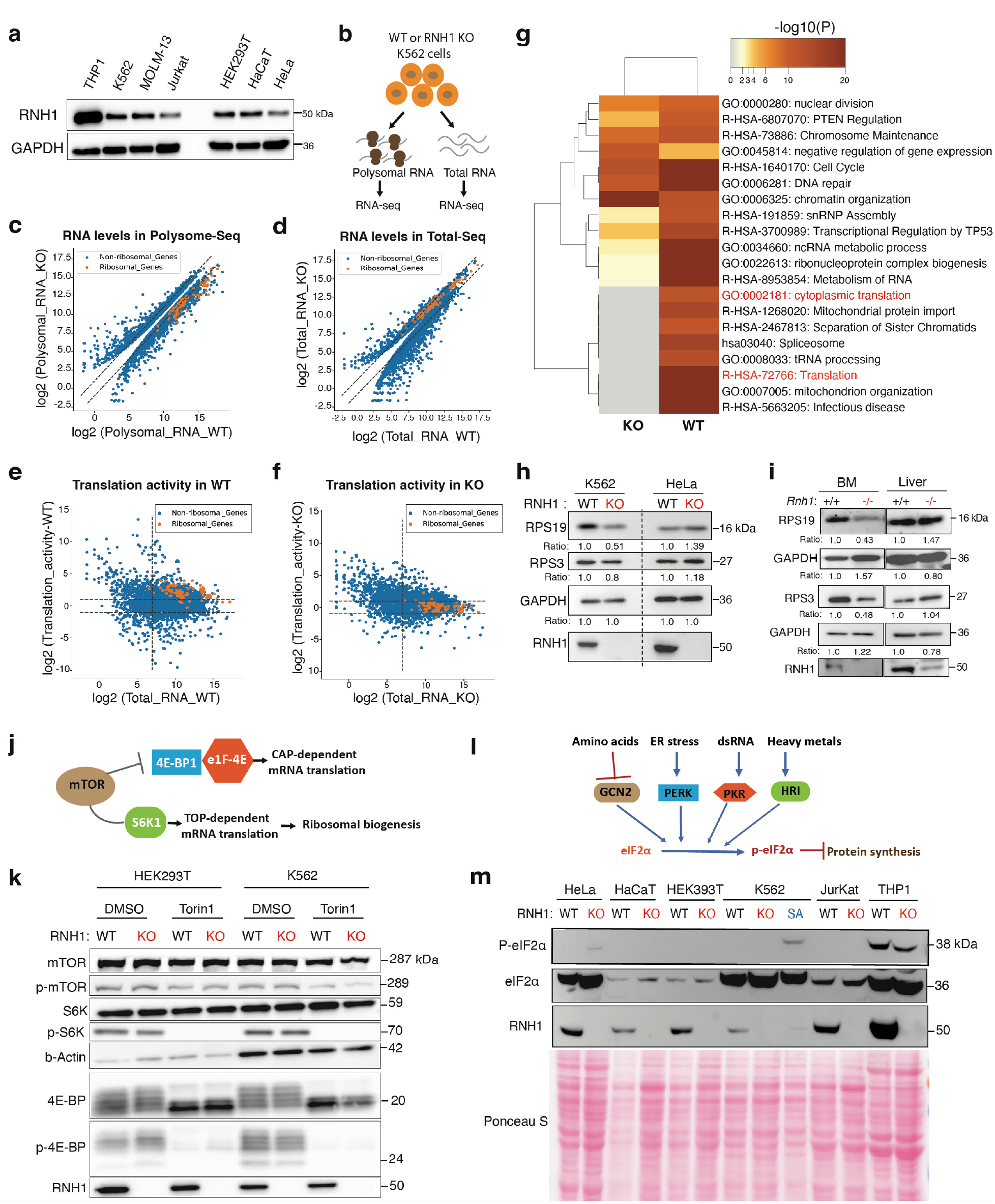
RNH1 is involved in ribosomal protein mRNA translation. (**a**) Total protein lysates from WT and RNH1 KO of hematopoietic and non-hematopoietic origin cell lines were analyzed by western blot with the indicated antibodies. Blots are representative of three independent experiments. (**b)** Schematic showing total and polysomal RNA isolation and RNA-seq analysis. (**c and d**) The expression of ribosomal and non-ribosomal genes in WT and RNH1 KO K562 cells obtained by polysomal RNA-seq (c) or total RNA-seq (d). Total genes with cutoff of padj. <0.1 in polysomal RNA-seq and total RNA-seq were selected and plotted. Dotted lines separate genes with log2(fold change) <1 and >1. (**e and f**) Plot of Log2 fold change of total RNA verses translational activity (TA) of ribosomal and non-ribosomal genes in WT (e) or RNH1 KO (f) samples. Horizontal dotted lines separate genes with log2(fold change) <1 and >1. Vertical line separate genes with arbitrary cutoff of >7< for log2(expression in total RNA-seq). (**g**) The heat maps generated via Metascape tool with default parameters show the relative expression of genes in different Gene ontology (GO) gene sets. (**h**) Protein lysates from K562 and HeLa with WT and RNH1 KO genotypes were analyzed by western blot for RPS19, RPS3 and RNH1. Blots are representative of three independent experiments. GAPDH is used as a normalizing control and numbers represent the normalized band intensity with respect to the corresponding WT sample. (**i**) Protein lysates from WT and *Rnh1^-/-^* bone marrow (BM) and liver were analyzed by western blot for RPS19, RPS3 and RNH1 (N=3 mice). GAPDH is used as a normalizing control and numbers represent the normalized band intensity with respect to the corresponding WT sample. Schematic of mTOR signalling. (**k**) WT and RNH1 KO HEK293T and K562 cells were treated with or without mTOR inhibitor Torin1 (100nM) and analyzed by western blot with the indicated antibodies. Blots are representative of four independent experiments. (**l**) Schematic of eIF2α-kinases mediated translation suppression. (**m**) WT and RNH1 KO HeLa, HACAT, HEK393T, K562, Jurkat and THP1 cell-lysates were analyzed by western blot with the indicated antibodies. Blots presented are representative of two independent experiments. K562 cells treated with sodium arsenate (SA) (100uM) used as positive control for stress induced translation inhibition.

To comprehend the mechanism by which RNH1 regulates RPs mRNAs translation, we investigated mTOR signaling in RNH1 KO and WT HEK293T and K562 cell lines. The mTOR pathway is known to regulate RPs mRNAs translation and global translation after nutrient sensing ^32^. However, we did not observe differences in the phosphorylation levels of mTOR, S6K and 4E-BP between WT and RNH1 KO cells in both cell-types (**Fig. 2j and k**). This suggests that RNH1 regulates RPs mRNA translation independent of mTOR signaling. Another important factor controlling global translation is eIF2α. Phosphorylation of eIF2α after cellular stress, mediated by eIF2α-kinases (GCN2, PERK, HRI and PKR) inhibits global translation and potentially contribute to the decrease in RPs mRNAs translation ^33^. However, we did not observe differences in the phosphorylation levels of eIF2α between WT and RNH1 KO cells in both cell-types (**Fig. 2l and m**). This excludes the possibility of stress-mediated translational defects in RNH1 KO hematopoietic cells. Collectively, these results indicate that mTOR signaling and eIF2α-phosphorylation, which are two major pathways involved in global translation regulation ^34^, do not account for the translational defects mediated by loss of RNH1 in hematopoietic cells.

### RNH1 binds to PABP and involved in mRNA circularization

Our previously published mass-spectrometry experiments have revealed that RNH1 directly binds to RPs and several RNA processing proteins ^21^. Among its interacting partners, poly(A) binding protein (PABP) caught our attention. Recently, another study also confirmed the binding of RNH1 to PABP ^35^. To further investigate these interactions, we performed co-immunoprecipitation of endogenous RNH1 interacting proteins using different cell lines (THP1, K562, and HEK293T) followed by mass-spectrometry (**Extended Data Fig. 5a and b and Supplementary Table. 3-5**). Supporting previous results, PABP was found to bind to RNH1 in all three cell lines (**Fig. 3a**). PABP is a well-known translation initiation factor that binds to the poly(A) tail of mRNAs ^36^. Its interaction with poly(A) tail stimulates translation by associating it with the eukaryotic initiation factor 4G (eIF4G) ^37^. This interaction helps to circularize mRNAs by bringing the 3′ end close to its 5′ m^7^G cap via sequential interactions of the 3′-poly(A)-PABP-eIF4G-eIF4E-5′ m^7^G cap ^36^. These interactions play pivotal roles in the translation initiation and maintaining translation efficiency in eukaryotic cells ^38^. Transcripts with a TOP (terminal oligopyrimidine tract) sequences, such as RPs, could be hypersensitive to defects in these interaction ^39^.

**Fig 3.**
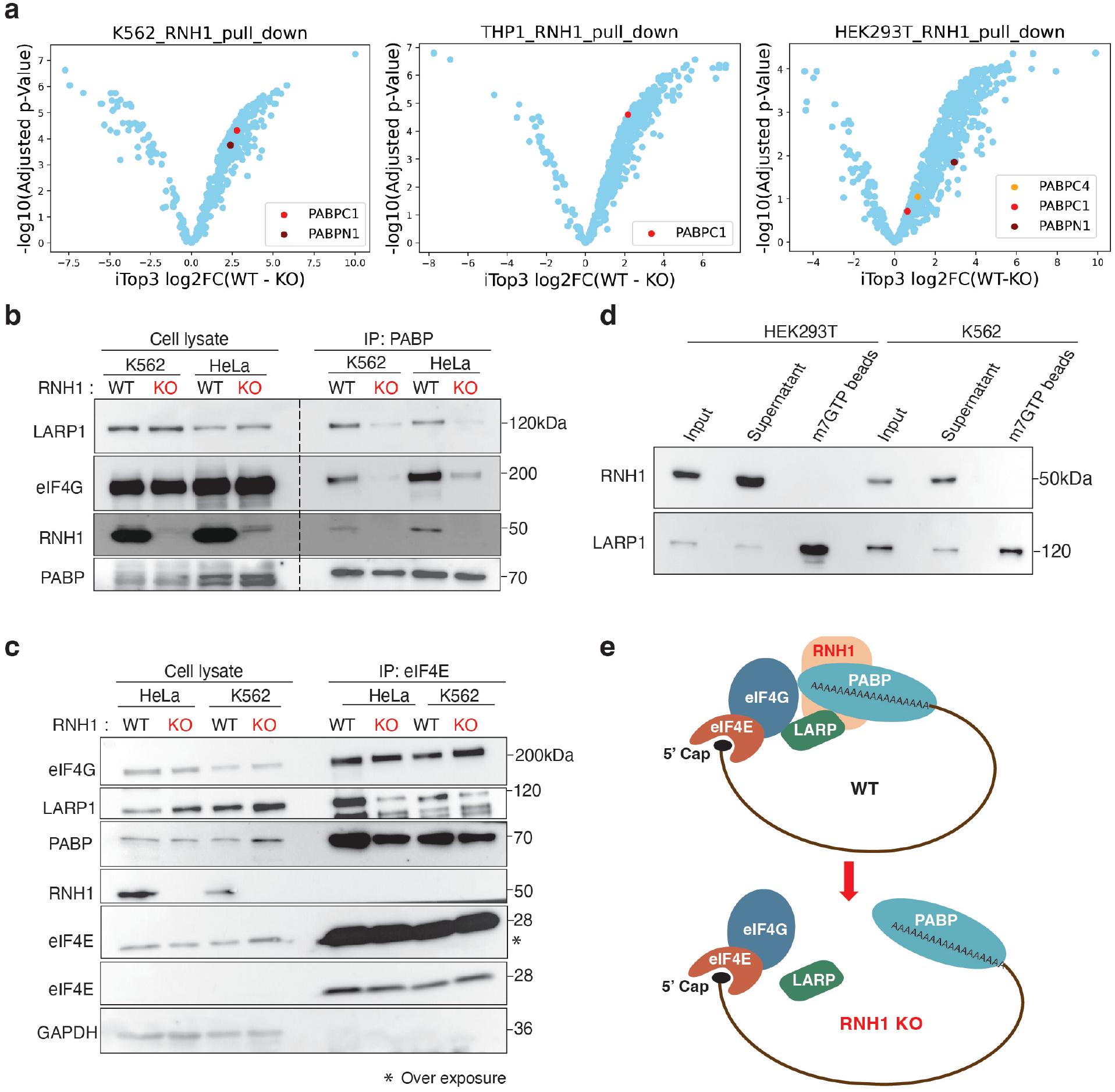
RNH1 bind to PABP and helps in mRNA circularization. (**a**) Volcano plots depicting RNH1 binding total proteome from WT K562, THP1 and HEK29T cells compared with RNH1 KO of corresponding cells. Plots show the comparisons of log2 fold changes (FC) versus adjusted p-values (FDR-controlled Benjamini and Hochberg multiple test correction) calculated based on peptide-imputed Top3 (iTop3). (**b, c**) Whole cell lysates of WT and RNH1 KO K562 and HeLa cells were used for anti-PABP immunoprecipitation (b) and anti-eIF4E immunoprecipitation (c), then immunoblotted, as indicated. Blots are representative of 3 independent experiments. (**d**) Whole cell lysates of K562 and HEK293T cells were used for m^7^GTP immunoprecipitation and immunoblotting, as indicated. Blots are representative of 3 independent experiments. (**e**) Schematics of RNH1 mediated mRNA circularization.

To check role of RNH1 in these interactions, PABP interacting partners from WT and RNH1 KO K562 cells were co-immunoprecipitated. As expected, eIF4G and RNH1 were immunoprecipitated in the PABP-IP of WT K562 cells (**Fig. 3b**). Surprisingly, PABP and eIF4G interactions were decreased in RNH1 KO K562 cells (**Fig. 3b**). This reduction in interaction between eIF4G⇔PABP might have prevented mRNA circularization and led to the observed translation decrease in RNH1 KO hematopoietic cells. LARP1 is another interacting partner of PABP involved in the translation of RPs mRNAs ^40^. Interestingly, LARP1 and PABP interaction also decreased in RNH1 KO K562 cells (**Fig. 3b**). The levels of PABP, eIF4G or LARP1 were not altered in WT and RNH1 KO K562 total cell lysates (**Fig. 3b**). This suggests the dysregulated assembly of factors at the 3’ end of mRNAs in the RNH1 KO K562 cells. To check the scenario at the 5’ end of mRNA, we pulled down eIF4E interacting partners, as eIF4E is a cap binding protein. Interestingly, the eIF4E-IP revealed that eIF4E-eIF4G interactions were not altered, but LARP1 and PABP interactions were reduced in RNH1 KO eIF4E-IP (**Fig. 3c**). This hints that assembly of eIF4E and eIF4G on 5’cap is unaltered in RNH1 KO cells. LARP1 and PABP were immunoprecipitated with eIF4E likely due to secondary interactions, as there are no direct interactions between LARP1⇔eIF4E⇔PABP known to our knowledge. Further, m^7^GTP cap binding assay revealed that RNH1 does not bind to the 5’cap in both hematopoietic and non-hematopoietic cells (**Fig. 3d**). Thus, RNH1 is absent at the 5’end and present at the 3’end of mRNAs via PABP, supporting reduced interactions between eIF4G⇔PABP but not between eIF4E⇔eIF4G (**Fig. 3b and c**). Thus, loss of RNH1 impact mRNA circularization and/or translation initiation via regulating LARP1⇔PABP⇔eIF4G interactions (**Fig. 3e**). To our surprise, LARP1⇔PABP⇔eIF4G interactions were also reduced in non-hematopoietic RNH1 KO HeLa cells, suggesting that RNH1 regulates LARP1⇔PABP⇔eIF4G interactions in both hematopoietic and non-hematopoietic cells (**Fig. 3b and c**). However, this raises the question of why the loss of RNH1 predominantly reduces translation in hematopoietic origin cells but not in non-hematopoietic origin cells.

### ANG compensates loss of RNH1 mediated translation defects in non-hematopoietic cells

To address the aforementioned discrepancy, we formulated a hypothesis that non-hematopoietic origin cells might possess specific compensatory mechanisms enabling them to tolerate the RNH1 loss. To explore potential candidates for further investigation, we sought out translation regulators that interact with RNH1 and are exclusively expressed in non-hematopoietic cells but not in hematopoietic origin cells. ANG emerged as a promising candidate, meeting all the criteria. RNH1 is known to bind to ANG and inhibit its functional activities ^13^. ANG exhibits a very weak ribonucleolytic activity but possesses strong capabilities in inducing blood vessel growth. It has been implicated in the initiation, progression, and metastasis of tumors ^13,41^. Furthermore, ANG plays contrasting roles in translation ^14,42^. In stress situations, it cleaves tRNA and decreases translation ^43^. Under favorable growth conditions, ANG translocates to the nucleus, triggering the synthesis of ribosomal RNA (rRNA), and actively participating in cell proliferation ^4,44^. To investigate the expression pattern of ANG in hematopoietic and non-hematopoietic origin primary human cells, we analyzed publicly available single-cell RNAseq data sets. Interestingly, we found minimal to no expression of ANG in hematopoietic cells compared to non-hematopoietic cells (**Fig. 4a**). In line with this, we did not find ANG protein expression in hematopoietic origin cell lines (**Fig. 4b**). This finding suggests that ANG might play a distinct role in translation regulation in non-hematopoietic cells, potentially contributing to their ability to cope with the loss of RNH1. The presence of ANG in the nucleus enhance rRNA synthesis ^4^, so we checked its localization by analyzing subcellular fractions of WT and RNH1 KO HeLa and K562 cells. As expected, ANG was not detected in both WT and RNH1 KO K562 cells (**Fig. 4c**). Surprisingly, in RNH1 KO HeLa cells, ANG expression increased in the nucleus and also other cell fractions compared to WT HeLa cells (**Fig. 4c**). Corroborating this, ANG levels were also increased in RNH1 KO HeLa and HaCaT total cell lysates (**Fig. 4d and e**). These results suggest that ANG is either expressed minimally or not at all in haemopoietic origin cells, but is expressed in non-hematopoietic origin cells. Additionally, the loss of RNH leads to increase in ANG expression and its translocation into the nucleus of non-hematopoietic origin cells.

**Fig 4.**
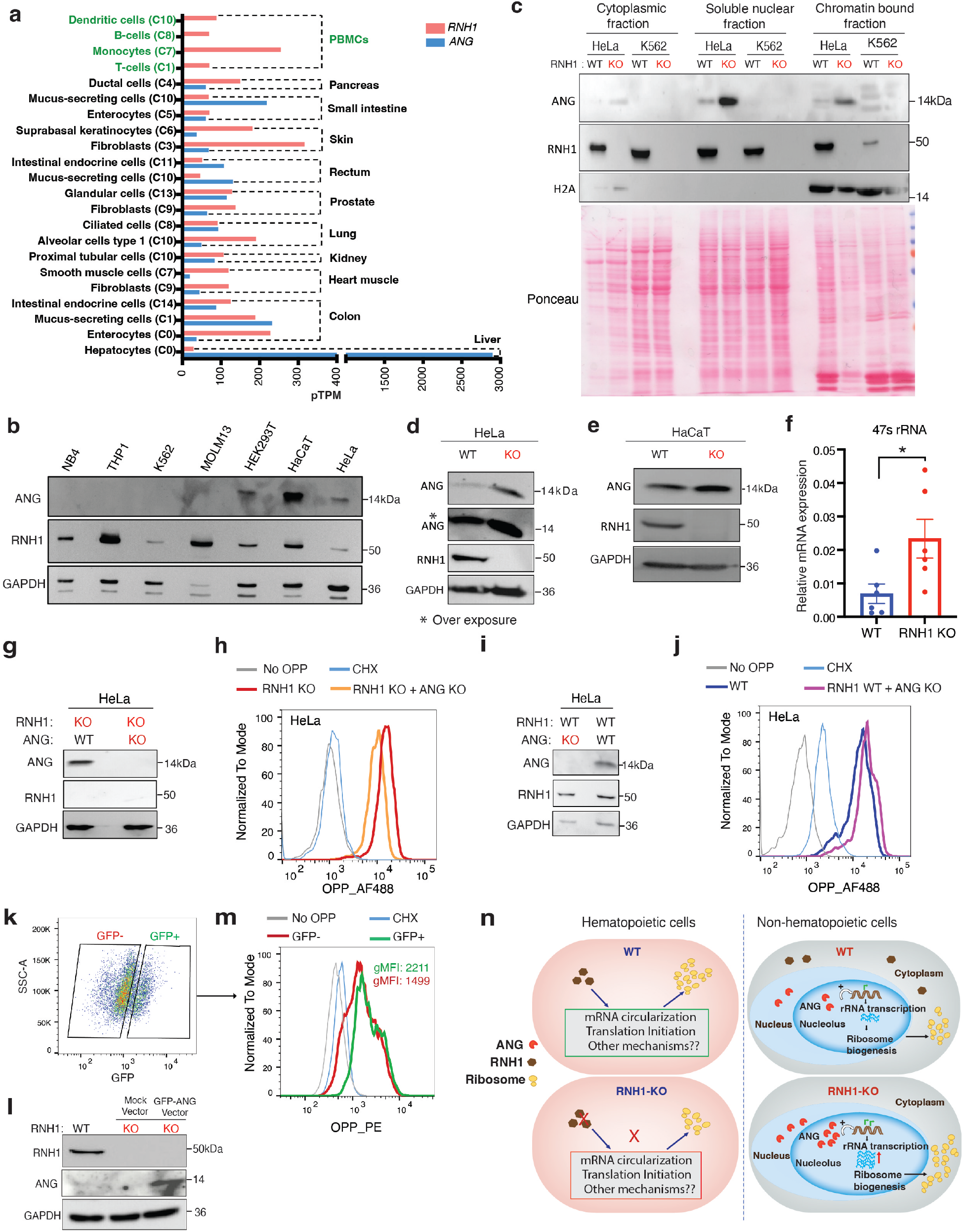
Angiogenin compensate translational defects in RNH1-deficient non-hematopoietic cells. (**a**) RNH1 and ANG mRNA expression from single-cell RNA sequencing of different human cells from hematopoietic and non-hematopoietic origin. c-0 to c-11 are cluster numbers. pTPM - protein-transcripts per million, data source: https://www.proteinatlas.org. (**b**) Total protein lysates from hematopoietic and non-hematopoietic origin cell line were analyzed by western blot with the indicated antibodies. Blots are representative of three independent experiments. (**c**) Total protein lysates of different cellular fractions from WT and RNH1 KO of K562 and HeLa cells were analyzed by western blot with the indicated antibodies. Blots are representative of three independent experiments. **(d and e**) Total protein lysates from WT and RNH1 KO HeLa (d) or HaCaT (e) cells were analyzed by western blot with the indicated antibodies. Blots are representative of three independent experiments. **(f**) qRT-PCR analysis of WT and RNH1 KO HeLa cells for 47S rRNA, normalized to 18S rRNA. Data are shown as mean ± SEM and are representative of 6 independent experiments. (**g**) Total protein lysates from RNH1 KO or RNH1 and ANG double KO HeLa cells were analyzed by western blot with the indicated antibodies. Blots are representative of three independent experiments. **(h**) RNH1 KO or RNH1 and ANG double KO HeLa cells were incubated for 1h with OPP and FACS analysis was performed to measure OPP incorporation. Representative histograms were shown for OPP fluorescence. Data is representative of 3 independent experiments. (**i**) Total protein lysates from WT or ANG KO HeLa cells were analyzed by western blot with the indicated antibodies. Blots are representative of three independent experiments. **(j**) WT or ANG KO HeLa cells were incubated for 1h with OPP and FACS analysis was performed to measure OPP incorporation. Representative histograms were shown for OPP fluorescence. Data is representative of 3 independent experiments. **(k-m**) RNH1 KO K562 cells were transduced with GFP-tagged ANG and incubated for 1h with OPP and FACS analysis was performed to measure OPP incorporation. FACS analysis shows GFP+ and GFP-expressing cells. Data is representative of 3 independent experiments (k). Total protein lysates from control or GFP-tagged ANG transduced RNH1 KO K562 cells were analyzed by western blot with the indicated antibodies. Blots are representative of three independent experiments (l). FACS analysis was performed to measure OPP incorporation in ANG-GFP transduced RNH1 KO K562 cells. Representative histograms were shown for OPP fluorescence (m). Data is representative of 3 independent experiments. gMFI (geometric mean fluorescence intensity). CHX (Cycloheximide). (**n**) Illustration of proposed RNH1/ ANG mediated cell-type-specific translational regulation.

To check whether the increased nuclear ANG enhance rRNA transcription in RNH1 KO cells (*41*), we quantified 47s rRNA levels in RNH1 KO HeLa cells via qPCR. Indeed, we found increased rRNA transcription in RNH1 KO HeLa cells (**Fig. 4f**), supporting the notion that ANG plays a compensatory role in these cells. These results indicate that ANG rescue the loss of RNH1 mediated translation defects in non-hematopoietic cells. This could explain the increased translation observed in RNH1 KO HeLa cells in luciferase reporter assay (**Fig. 1d**) as well as the elevated RP expression (**Fig. 2h**). To further ascertain that ANG mediates compensatory mechanism, we deleted ANG in RNH1 KO HeLa cells and monitored translation by OPP incorporation (**Fig. 4g**). As hypothesized, loss of ANG decreased translation in RNH1 KO HeLa cells. (**Fig. 4h)**, while ANG deletion in WT HeLa cells had no impact on translation (**Fig. 4i and j**). These results indicate that the loss of ANG has no impact on translation under steady-state conditions. Additionally, we investigated whether expressing ANG in RNH1 KO K562 cells could rescue translational defects. We ectopically expressed GFP-tagged ANG in RNH1 KO K562 cells (**Fig. 4k and l**) and monitored translation by OPP incorporation. Supporting the above results, ANG-expressing RNH1 KO K562 cells (GFP+) showed increased translation compared to RNH1 KO K562 cells without ANG expression (GFP-) (**Fig. 4m**). Collectively, these results suggest that ANG-mediated translation compensates for the translation defects caused by the loss of RNH1 in non-hematopoietic cells (**Fig. 4n**). This compensatory mechanism involving ANG could be crucial for maintaining translation efficiency and cellular functions in the absence of RNH1.

We next asked whether RNH1 mediated cell-type-specific translational defects result in tissue-specific functional impairments. Our previous findings demonstrated that the loss of RNH1 leads to defects in embryonic erythropoiesis. Further, knockdown of RNH1 in human CD34^+^ cells decreased erythroid differentiation ^21^. Similarly, deletion of RNH1 in mouse hematopoietic cells (HSCs) was found to dysregulate hematopoiesis and cause anemia ^45^. Moreover, mutations in the RNH1 gene in human have also been associated with anemia phenotype ^22^. Overall, these studies collectively provide compelling evidence that the loss of RNH1 leads to defects or dysregulation in hematopoiesis. Interestingly, in contrast to the hematopoietic system, we observed no major phenotypic differences in liver-specific *Rnh1^-/-^*(*Rnh1^fl/fl^, Alb-Cre^Tg/+^*) mice compared to WT. Liver-specific *Rnh1^-/-^* mice exhibited normal survival rates, and there were no changes in body as well as liver weight (**Extended Data Fig. 6a-c**). Additionally, liver function tests, such as alanine transaminase (ALT) and aspartate transaminase (AST) assays, also showed no significant differences between WT and liver-specific *Rnh1^-/-^* mice (**Extended Data Fig. 6d and e**). Furthermore, blood glucose and peripheral blood levels were also similar in both groups (**Extended Data Fig. 6f and g**). Moreover, we analyzed these mice for 18 months and did not notice any phenotypic differences between WT and liver-specific *Rnh1^-/-^* mice (data not shown). Although we cannot completely exclude the possibility of translation-independent RNH1 functions contributing to tissue-specific phenotype, the absence of major phenotypic differences in liver-specific *Rnh1^-/-^* mice strongly indicate the absence of RNH leads to tissue-specific defects, consistent with the observed translation defects in specific tissues.

Collectively, these findings unveil the existence of cell-type-specific global translation regulation, with RNH1, a ribosome-associated protein, emerging as a key regulator of translation. Our results suggest that RNH1 potentially governs the global translation by controlling the translation of ribosomal protein transcripts through regulating mRNA circularization. Nevertheless, further investigations are required to fully comprehend the precise mechanisms by which RNH1 mediates translation regulation. Consistent with the role of RNH1 in translation, we observed a decrease in translation upon the loss of RNH1 in hematopoietic-origin cells. However, this decrease in translation was not observed in non-hematopoietic origin cells. Our results conclusively demonstrate that this translation specificity is mediated by a compensatory mechanism regulated by ANG. In non-hematopoietic origin cells, ANG is expressed, and the loss of RNH1 further amplifies its expression. This increased ANG expression enhances translation through rRNA synthesis and compensates for the translation defects caused by RNH1 loss. Conversely, hematopoietic-origin cells either do not express ANG or express it at very low levels, making them susceptible to translational defects following RNH1 loss (**Fig. 4n**). While our results strongly indicate that the loss of RNH1 significantly reduces translation in hematopoietic origin cells, we cannot dismiss the possibility that other cell-types might also be vulnerable to RNH1-mediated translation defects, considering the existence of over 200 different cell-types in the human body Genes involved in translation, such as ribosomal protein genes, are highly conserved across eukaryotes. However, RNH1 and ANG gene have only evolved in vertebrates (**Extended Data Fig. 1**) ^47^, suggesting that vertebrates have developed an additional layer of translation control through cell-type-specific regulation. These regulatory or compensatory mechanisms may play a crucial role in facilitating efficient protein translation and ensuring overall organismal survival. Although further investigation is warranted, it is tempting to speculate that these compensatory mechanisms might contribute to the specific phenotypes observed in ribosomopathies. Our study serves as a prime example of the regulation of cell-type-specific global translation, highlighting the significance of comprehending these regulatory mechanisms to gain a comprehensive understanding of the translation process in vertebrates. Considering the significant role of translation dysregulation in cancer progression, neurological diseases, and ribosomopathies ^6,34,48,49^, understanding tissue-specific translation regulators would be highly advantageous for targeting these specific diseases.

## Methods

### Generation of mouse model and *in vivo* OPP incorporation studies

Generation of *Rnh1* conditional-knockout (*Rnh1^fl/fl^*) mice were reported previously ^16^. We crossed *Rnh1^fl/fl^* mice with *Mx1*-*Cre* (Jackson: 002527) and Albumin Cre mice (Jackson: 0103574) to generate inducible *Rnh1^-/-^* mice (*Rnh1^fl/fl^ Mx1-Cre**^+^***) and liver specific *Rnh1^-/-^* mice (*Rnh1^fl/fl^, Alb-Cre^Tg/+^*), respectively. *Rnh1* was excised in *Rnh1^fl/fl^ Mx1-Cre**^+^*** mouse model by giving three rounds of 200 μg poly(I:C) (Invivogen) using intraperitoneal injections once every two days. The screening for *Rnh1* deletion after Cre-recombination was performed by PCR genotyping as described previously ^16^. Western blot was also performed using spleen and liver to check RNH1 protein expression. The genotypes of the mice could not be blinded or randomized, due to the experimental design. All groups of mice (experimental and control mice) were age- and sex-matched. We used littermates for our experiments. Mice were maintained at SPF (Specific Pathogen Free) conditions at University Bern, Switzerland. All animal experiments were approved by the Swiss Federal Veterinary Office (Bern, Switzerland) under valid authorization (BE19/2019, BE103/2022). Mice were handled according to Swiss Federal Veterinary Office guidelines under valid authorization.

*Rnh1^-/-^* mice (*Rnh1^fl/fl^ Mx1-Cre**^+^*** and *Rnh1^fl/fl^, Alb-Cre^Tg/+^* mice) and wildtype littermates (*Rnh1^fl/fl^ Cre**^-^***) were injected intraperitoneally with O-propargyl-puromycin (OPP) (Invitrogen) (50mg/kg). PBS was injected in the vehicle group. After one hour, the organs were collected and embedded in paraffin. Staining was performed as described previously ^28^. Briefly, after fixation and permeabilization, tissue slides were stained by copper(I)-catalyzed azide-alkyne cycloaddition (CuAAC) with the tetramethylrhodamine (TMR)-azide (Click it reaction) to fluorescently label OPP incorporated peptides. Finally, the tissue sections were counterstained with the Hoechst, mounted in standard mounting media, and then imaged using Nikon Ti-E fluorescence microscope.

### Cells and cell culture media

Cell lines K562, HEK293T, THP1, MOLM13, Jurkat, HeLa, HaCaT, and SHSY5Y were grown in RPMI 1640 GlutaMAX™-I medium (Invitrogen) or DMEM medium (500 mL, ThermoFisher) supplemented with 10% (vol/vol) FBS (Amimed) and 1% of penicillin and streptomycin (PAA Laboratories) and incubated at 37°C and 5% CO2. K562, HEK293T, THP1, Jurkat, HeLa and HaCat cell lines were described previously ^21,50,51^. Cell lines were tested negative for mycoplasma contamination using MycoAlert Mycoplasma Detection Kit (Lonza, Cat#LT07-318). MOLM13 cells were purchased from ATCC. Jurkat, HeLa cells were kindly provided by P. Schneider (University of Lausanne, Lausanne, Switzerland), HaCatT cells were kindly provided by E.J. Müller (University of Bern, Bern, Switzerland). SH-SY5Y cells were kindly provided S. Saxena (University of Bern, Bern Switzerland).

### Reagents and plasmids

We purchased cycloheximide (CHX), sodium arsenite from Sigma-Aldrich, mTOR inhibitor Torin-1 from Invivogen, Click-iT reaction kit and dynabeads from Invitrogen (Thermoscientific). Full-length RNH1-Flag construct and full length GFP-RNH1 plasmids were described previously^21^. To generate Full length GFP-ANG plasmid, the human Angiogenin (ANG) full coding sequence (ORIGEN RC 208874) was cloned into the entry vector pENTR4-GFP-C1 (W392-1, Addgene) and then recombined into the destination vector pLenti CMV Blast DEST (706-1, Addgene) plasmid using the Gateway cloning method.

### Generation of CRISPR/CAS9-mediated RNH1 and ANG knockout cells

RNH1 KO cell lines were generated as described previously ^21^. For generation of ANG KO in WT and RNH1-KO HeLa cells, CRISPR gRNA sequences targeting exon 1 on different regions of Human ANG were designed and KO cells were obtained as described previously ^21^. Target exon and the seed sequences preceding the protospacer adjacent motif (PAM) are as follows: ANG_sgRNA1_F: 5’-CACCGAACGTTTCTGAACCCCGCTGTGG-3’, ANG_sgRNA1_R: 5’AAACCCACAGCGGTTCAGAAACGTTC3’. Primer2: ANG_sgRNA2_F: 5’CACCGTGGCAACAAGCGCAGCATCAAGG3’, ANG_sgRNA2_R:5’AAACCCTTGA TGCTGCGCTTGTTGCCAC 3’. All generated knockout clones were tested negative for mycoplasma contamination using MycoAlert Mycoplasma Detection Kit (Lonza, Cat#LT07-318).

### Reconstitution of Flag-RNH1 in RNH1-KO K562 cells

Flag-RNH1 was sub-cloned into the retroviral vector pMSCVpuro (Clontech) as described previously ^21^. Retroviral vector pMSCVpuro-Flag–RNH1 was co-transfected with the helper plasmids VSV-G and Hit60 into HEK293T cells using PEI transfection reagent. Culture supernatants containing recombinant viral particles were harvested and used to infect RNH1-KO K562 cells. To establish stable cell lines, puromycin (5 µg/mL) was added to the cells three days after infection for selection.

### Generation of RNH1-KO K562 cells expressing GFP-ANG

RNH1-KO K562 cells were infected with lentiviruses expressing ANG (pLenti CMV Blast GFP-ANG) or the empty vector pLenti CMV Blast DEST (706-1, Addgene) as previously described ^52^. After 48 hours post-infection, cells were used for OPP incorporation experiments.

### In vitro OPP and AHA incorporation assays

To measure the translation in cells, we performed O-propargyl Puromycin (OPP) incorporation assays. OPP is an analog of puromycin that gets incorporated into nascent polypeptides. Cells were incubated with OPP (50 uM) for one hour and then washed with PBS. Click-iT reaction was performed using Alexa Fluor 488 (AF488)-azide and the Click-iT Protein Reaction Buffer Kit (C10276, Invitrogen) for 30 min according to the manufacturer’s protocol.

For cell lines resistant to puromycin, we used AHA (l-azidohomoalanine) instead of OPP. To deplete methionine reserves, cells were incubated for an hour in a methionine-free RPMI medium (R7513, Sigma-Aldrich) containing 2 mM l-glutamine. Then, cells were washed and treated with AHA (C10102, Life Technologies) at a final concentration of 50 μM for one hour. After incubation, cells were washed with PBS, and Click-iT reaction was performed using TAMRA alkyne and the Click-iT Protein Reaction Buffer Kit (C10276, Invitrogen) for 30min according to the manufacturer’s protocol.

Hepatocyte isolation

Hepatocytes were isolated from wildtype (*Rnh1^fl/fl^*) and *Rnh1^-/-^* mice (*Rnh1^fl/fl^, Alb-Cre^Tg/+^*) in situ using a two-step collagenase perfusion method, as previously described ^53^. The isolated live hepatocytes were plated and subjected to OPP incorporation assays as described above.

### Luciferase reporter assays

WT or RNH1 KO cells were transfected with plasmid containing luciferase reporter gene having b-actin or GAPDH 5’UTR ^3^. Lipofectamine 3000 used for transfection as per manufacturer’s instructions. After 48 hours of incubation, cells were washed and lysed in a lysis buffer (E1500 Promega). Lysates were mixed with Firefly luciferase assay reagent (E1500 Promega) as per manufacturer’s protocol and scanned for luminescence in Tecan scanner.

### Polysome profile analysis and fractionation

Polysome profiling was performed as described previously ^21^. Briefly, 5-8 × 10^6^ cells were washed with PBS containing cycloheximide (CHX) (100 ug/ml) and resuspended in 200 μl of hypotonic buffer (1.5 mM KCl, 2.5 mM MgCl2, and 5.0 mM Tris-Cl, pH 7.4) and 200 μl hypotonic lysis buffer (same with 2% sodium deoxycholate, 2% Triton X-100, and 2.5 mM DTT) plus CHX and gently disrupted using a Dounce homogenizer. The lysates were centrifuged at 8,000 g for 10 minutes at 4°C. The optical density of supernatant sample was measured on an Ultrospec UV/Visible spectrophotometer 1000 (Pharmacia Biotech). Equal OD of the supernatant was supplemented with 80 μl heparin. Linear 10% to 50% sucrose gradients (80 mM NaCl, 5 mM MgCl2, 20 mM Tris-Cl, pH 7.4, and 1 mM DTT) were prepared manually. The supernatant was loaded onto the pre-prepared sucrose gradients. Gradients were placed onto a pre-cooled SW41 Ti rotor (Beckman Coulter) and were centrifuged at 39,000 rpm for 2.45 hours at 4°C using Optima XPN-80 Ultracentrifuge (Beckman Coulter). Following centrifugation, gradients were replaced onto a gradient density fractionation system (Brandel) and separated through a live OD 254 nm using UA-6 UV/Vis detector (Teledyne Isco). Polysome fractions were collect using Foxy R1 fraction collector (Teledyne Isco).

### RNA isolation and qRT-PCR

Total RNA was isolated from cells using the QIAGEN RNeasy Kit according to the manufacturer’s protocol. Polysomal RNA was isolated from polysome fractions by TrizolTM (Invitrogen) according to the manufacturer’s protocol. Reverse transcription and real-time PCR (RT-PCR) from total RNA was carried out as described previously ^21^. The SYBR Green Dye detection system was used for quantitative real-time PCR on Light Cycler 480 (Roche). Gene-specific primers (Microsynth) used are listed in **Table. S6**. Reactions containing of ddH2O instead of cDNA were used as negative controls for target and housekeeping genes.

### tiRNA gel electrophoresis

HeLa and K562 cells were cultured in a 6 well plates. For positive control, cells were treated with 250 uM sodium arsenite for 2 hrs. Total RNA was isolated from all samples as described above and heated with RNA loading dye at 60^°^ C for 5 min. Then, RNA separated on 15% urea PAGE gel and tiRNAs detected as described previously ^54^.

### RNA seq experiments and Bioinformatic data analysis

RNA-sequencing from total RNA was reported previously^21^. For Polysome RNA sequencing, between 200 ng to 1μg of high-quality RNA (RIN > 7) samples were used for cDNA synthesis and library preparation using the TruSeq Stranded total RNA Sample Preparation kit (Illumina). The cDNA libraries were then sequenced with 2 × 150 bp reads on an Illumina HiSeq3000 instrument. The quality of the RNA-seq data was assessed using fastqc v. 0.11.2 (Babraham Bioinformatics), RSeQC v. 2.6.1^55^, and Qualimap v. 2.2.1^56^. The readings were mapped to the human reference genome (assembly GRCh38) using HiSat2 v. 2.1.0 ^57^. We then used Feature Counts from the Subread v. 1.5.3 ^58^ to count the number of readings overlapping with each gene as specified in the Ensembl annotation. For each total RNA and polysomes RNA-seq data, the Bioconductor package DESeq2 v. 1.18.0 ^59^ was used separately to test for the differential gene expression between RNH1-deficient cells and controls. The TopGo was used to identify gene ontology terms containing unusually many differentially expressed genes^60^. From the above gene expression file, total genes (4029) with a cutoff of padj <0.1 in Polysomal RNA-seq and Total-seq were selected. This includes the expression of ribosomal (152) and non-ribosomal genes (3877) in wildtype (WT) and RNH1-deficient (KO) K562 cells. The values of Log2(expression in total RNA-seq of WT) were plotted against the Log2 (expression in total RNA-seq of KO) using custom Python script. Similarly, the values of Log2(expression in polysome RNA-seq of WT) were plotted against the Log2(expression in polysome RNA-seq of KO). Dotted lines separate genes with log2(fold change) <1 and >1. Joint analysis of the polysomal and the total RNA was performed with anota2seq v. 1.0.1 in R v. 3.4.2 to assign genes to different regulatory modes^31^. Anota filtered genes (262) were plotted as described in the above paragraph.

To find the change in the translational activity of genes, 4029 genes with a cutoff of padj. <0.1 in Polysomal RNA-seq and Total RNA-seq were selected. The translation activity (TA) of these genes were calculated as polysomal RNA/ Total RNA. The values of log2(TA) were plotted with log2(Total RNA). Dotted lines separate genes with log2(fold change) <1 and >1. The translational activity of Anota filtered genes (262) were also calculated and plotted as described in the above paragraph.

### **I**mmunoblot analysis

Cells from different cell lines, mouse total BM cells and hepatocytes were resuspended in the lysis buffer (20 mM Tris, pH 7.4, 150 mM NaCl, 1% (vol/vol) Nonidet-P40, 10 mM EDTA). Extracts were used for immunoblot after removing debris. Nitrocellulose membranes were stained with Ponceau S to ensure equal protein loading. The following antibodies were used for immunoblots.

| Antibody | Reactivity | Manufacturer | Reference |
| --- | --- | --- | --- |
| anti-RNH1 (A-9) | Human | Santa Cruz Biotechnology | sc-365783 |
| anti-RNH1 (C-10) | Mouse | Santa Cruz Biotechnology | sc-271725 |
| anti-GAPDH (G-9) | Human, Mouse | Santa Cruz Biotechnology | sc-365062 |
| anti-β-actin | Human, Mouse | Abcam | ab8227 |
| anti- ANG | Human | Santa Cruz Biotechnology | sc-74528 |
| anti-mTOR (7C19) | Human | Cell Signaling | 2983T |
| anti-P-mTOR (Ser2448) | Human | Cell Signaling | 5536T |
| anti-P-4E-BP1 (Thr37/46) | Human | Cell Signaling | 2855T |
| anti-4-EB-P1 | Human | Cell Signaling | 9644T |
| anti-p70 S6Kinase (Thr389) | Human | Cell Signaling | 9234T |
| anti-p70 S6 Kinase | Human | Cell Signaling | sc-8418 |
| anti RPL11 | Human, Mouse | Santa Cruz Biotechnology | sc-25931 |
| anti-RPS3 | Human, Mouse | Santa Cruz Biotechnology | sc-376008 |
| anti-RPS21 | Human, Mouse | Santa Cruz Biotechnology | sc-514411 |
| anti-H2A | Human | Cell Signaling | 12349T |
| anti-RPS19 | Human, Mouse | Santa Cruz Biotechnology | sc-100836 |
| anti-PABP1 | Human | Santa Cruz Biotechnology | sc-32318 |
| anti-eIF4E | Human | Santa Cruz Biotechnology | sc-9976 |
| anti-eIF4G | Human | Cell Signaling | 2469T |
| anti-LARP1 | Human | Santa Cruz Biotechnology | sc-515873 |
| anti-eIF2a | Human, Mouse | Santa Cruz Biotechnology | sc-133132 |
| anti-P-eIF2a | Human, Mouse | Cell Signaling | 9721 |

### Co-Immunoprecipitation experiments

Wild type or RNH1-KO cells were resuspended in the lysis buffer (20 mM Tris, pH 7.4, 100 mM NaCl, 1% (vol/vol) Triton x 100, 10 mM EDTA. Dynabeads were incubated with anti-PABP1 or anti-eIF4E antibody for 20 min. and then washed with PBS + tween 20. Subsequently, antibody bound dynabeads were incubated overnight with the cell lysate at 4° C. After washing, beads were resuspended in sample buffer and analyzed via immunoblot.

### Cap-binding assay

For cap-binding assay, cell lysates were prepared as described above. Then, 50 μl of the blank agarose bead control slurry and 50 μl of the γ-aminophenyl-m7GTP agarose C10-linked bead slurry (Jena Biosciences) were taken to separate 1.5 ml microcentrifuge tubes, centrifuged, and washed. Cell lysates were incubated with blank beads for 10min at 4° C. Then, this mixture was centrifuged, and supernatant was transferred to the tube containing γ-aminophenyl-m7GTP agarose C10-linked beads. Beads and lysates were incubated together overnight at 4° C on a rotor. Then, the beads were washed, resuspended in sample buffer, and analyzed via immunoblot.

### Mass spectrometry analyses

Wild type or RNH1-KO human THP1, K562, HEK293T cells (5x10^6^) were resuspended in the lysis buffer (20 mM Tris, pH 7.4, 150 mM NaCl, 1% (vol/vol) Nonidet-P40, 10 mM EDTA. Extracts were immunoprecipitated with anti-RNH1 magnetic beads and then were assessed by immunoblot. For mass spectrometry analyses, beads after immunoprecipitation were washed with PBS and resuspended in 30 μL 8 M urea, 50 mM Tris/HCl pH 8 and proteins reduced (10 mM DTT, at 37°C for 30 min), followed by alkylation with 50 mM iodoacetamide (at 37°C for 30 min in the dark) and digestion by 100 ng trypsin overnight at room temperature after dilution of urea to 1.3 M with 20 mM Tris/HCl, 2 mM calcium di-chloride pH 8.0. The digests were acidified with TFA and analyzed by nano-liquid chromatography (LC)-MS/MS (PROXEON coupled to a QExactive HF mass spectrometer, ThermoFisher Scientific) with 3 injections of 5 μl digest each. Peptides were trapped on a micro-precolumn C18 PepMap100 (5μm, 100 Å, 300 μm×5mm, ThermoFisher Scientific, Reinach, Switzerland) and separated by backflush onto the analytical nano-column (C18, 3µm, 155Å, 0.075 mm i.d. x 150mm length, Nikkyo Technos, Tokyo, Japan) applying a 40-minute gradient of 5% acetonitrile to 40% in water, 0.1% formic acid, at a flow rate of 350 nl/min. The Full Scan method was set with resolution at 60,000 with an automatic gain control (AGC) target of 1E06 and maximum ion injection time of 50 ms. The data-dependent method for precursor ion fragmentation was applied with the following settings: resolution 15,000, AGC of 1E05, maximum ion time of 110 milliseconds, mass window 1.6 *m*/*z*, collision energy 27, under fill ratio 1%, charge exclusion of unassigned and 1+ ions, peptide match preferred, and dynamic exclusion of fragmented precursors for 20 sec., respectively.

Mass spectrometry data was interpreted with MaxQuant (version 1.5.4.1) ^61^ against a forward and reversed Swissprot human protein sequence fasta file (release December 2017) using the following settings: strict trypsin cleavage rule allowing up to three missed cleavage sites, variable modifications of acetylated protein N-termini and oxidation of methionine and fixed carbamidomethylation of cysteines; peptide and fragment mass tolerances of 10 and 20 ppm; match between runs of the same sample type activated, but not between different types, in order to avoid over-interpretation. Identified peptides and proteins were filtered to a 1 % false discovery rate based on reversed database matches and a minimum of two razor or unique peptides were requested for acceptance of protein group identification.

### Phylogenetic tree reconstruction

Human RNH1 DNA sequence (NM_203387.3) was used as a query to perform nucleotide Blast with selected species. From Blast output, only those pairs that had query cover more than 70% (except for non-mammalian species) and percent identity more than 60 were selected. No significant similarity found below class “Reptilia” with above criteria. DNA sequences of selected species with these criteria were aligned using MAFFT multiple sequence aligner^62^. Phylogenetic analysis was performed on IQ-Tree webserver (http://iqtree.cibiv.univie.ac.at/) using ultrafast bootstrap analysis with 1000 number of bootstrap alignments. Phylogenetic tree was visualized using Interactive Tree Of Life (iTOL). Sequences used for multiple sequence alignment are: NM_203387.3 *Homo Sapiens;* XM_024241348.1 *Pongo abelii;* XM_004050348.3 *Gorilla gorilla gorilla;* NM_001009060.1 *Pan troglodytes;* XM_001488475.4 *Equus caballus;* NM_145135.3 *Mus musculus;* NM_001047113.1 *Rattus norvegicus;* NM_001006473.1 *Gallus gallus;* XM_019485196.1 *Alligator mississippiensis*, XM_008120593.1 *Anolis carolinensis*

### Statistical analysis

Data were expressed as mean ± SEM of at least three biological replicates or three independent replicates from each genotype. Comparison between two groups was performed by two-tailed *t*-test. A value of p<0.05 was considered to be statistically significant. All statistical analyses were calculated using Graph Pad Prism V. 9.5.1. No statistical methods were used to predetermine sample size. The experiments were not randomized and the investigators were not blinded to allocation during experiments and outcome assessment. Polysome RNAseq experiments were performed using three independent replicates from each genotype. Mass spectrometry studies were also performed on three independent replicates.

### Data and materials availability

All datasets generated and analysed during the current study are available in Tables. S1 to S5. Polysome RNA-seq data that support the findings of the study have been deposited at GEO under accession number GSE237169. The total RNA-seq data used in this study was deposited and accessed from the European Nucleotide Archive (ENA Data https://www.ebi.ac.uk/ena/) under accession number PRJEB35613.

## Acknowledgments

We thank lab members Christoph Liechti and Julia Aebischer for their assistance in some experiments. We thank Vijay Sankaran (Harvard medical school, USA) for Actin and GAPDH luciferase plasmids. We thank FACS, mass spectrometry and microscopy facility of the DBMR (University of Bern) for assistance in flow cytometry, mass spec and fluorescence microscopy experiments, respectively. We thank the Next Generation Sequencing (NGS) Platform (University of Bern) for RNA-Seq studies and Interfaculty Bioinformatics Unit for bioinformatic analysis. This work was supported by the Swiss National Science Foundation (PP00P3_157486, PP00P3_183721 & PP00P3_190073) to RA.

## Author Contributions

MAS and MS performed most of the experimental work and analysed the data. IK helped with RNA seq data analysis. AT maintained mouse lines and helped with experimental work. NA, FG, GB, CA helped with experimental work. AK helped experiments with hepatocytes. AAS helped with reagents, methodology and advice. M. Heller helped with mass spec data. MAS and RA wrote the manuscript with inputs from all the authors. RA conceived, acquired the funding, and supervised the research work.

## Competing interests

None to declare.

## Notes

### Competing Interest Statement

The authors have declared no competing interest.

## References

1. Xue, S. & Barna, M. Specialized ribosomes: a new frontier in gene regulation and organismal biology. Nat. Rev. Mol. Cell Biol. 13, 355–369 (2012).

2. Kondrashov, N. et al. Ribosome-mediated specificity in Hox mRNA translation and vertebrate tissue patterning. Cell 145, 383–397 (2011).

3. Ludwig, L. S. et al. Altered translation of GATA1 in Diamond-Blackfan anemia. Nat Med 20, 748–753 (2014).

4. Tsuji, T. et al. Angiogenin is translocated to the nucleus of HeLa cells and is involved in ribosomal RNA transcription and cell proliferation. Cancer Res. 65, 1352–1360 (2005).

5. Harbison, C. T. et al. Transcriptional regulatory code of a eukaryotic genome. Nature 431, 99– 104 (2004).

6. Narla, A. & Ebert, B. L. Ribosomopathies: human disorders of ribosome dysfunction. Blood 115, 3196–3205 (2010).

7. Horos, R. & Lindern, M. von. Molecular mechanisms of pathology and treatment in Diamond Blackfan Anaemia. Br J Haematol 159, 514–27 (2012).

8. Horos, R. et al. Ribosomal deficiencies in Diamond-Blackfan anemia impair translation of transcripts essential for differentiation of murine and human erythroblasts. Blood 119, 262–272 (2012).

9. Genuth, N. R. & Barna, M. Heterogeneity and specialized functions of translation machinery: from genes to organisms. Nat. Rev. Genet. 1–22 (2018) doi:10.1038/s41576-018-0008-z.

10. Mills, E. W. & Green, R. Ribosomopathies: There’s strength in numbers. Science 358, eaan2755 (2017).

11. Dutt, S. et al. Haploinsufficiency for ribosomal protein genes causes selective activation of p53 in human erythroid progenitor cells. Blood 117, 2567–2576 (2011).

12. Warner, J. R. & McIntosh, K. B. How Common Are Extraribosomal Functions of Ribosomal Proteins? Mol. Cell 34, 3–11 (2009).

13. Dickson, K. A., Haigis, M. C. & Raines, R. T. Ribonuclease inhibitor: structure and function. Prog. Nucleic Acid Res. Mol. Biol. 80, 349–374 (2005).

14. Sarangdhar, M. A. & Allam, R. Angiogenin (ANG)—Ribonuclease Inhibitor (RNH1) System in Protein Synthesis and Disease. Int J Mol Sci 22, 1287 (2021).

15. Haigis, M. C., Haag, E. S. & Raines, R. T. Evolution of ribonuclease inhibitor by exon duplication. Mol. Biol. Evol. 19, 959–963 (2002).

16. Bombaci, G. et al. LRR-protein RNH1 dampens the inflammasome activation and is associated with COVID-19 severity. Life Sci Alliance 5, e202101226 (2022).

17. Tavernier, Q. et al. Regulation of IRE1 RNase activity by the Ribonuclease inhibitor 1 (RNH1). Cell Cycle 17, 1901–1916 (2018).

18. Yao, X. et al. A novel role of ribonuclease inhibitor in regulation of epithelial-to-mesenchymal transition and ILK signaling pathway in bladder cancer cells. Cell Tissue Res. 353, 409–423 (2013).

19. Yamasaki, S., Ivanov, P., Hu, G.-F. & Anderson, P. Angiogenin cleaves tRNA and promotes stress-induced translational repression. J. Cell Biol. 185, 35–42 (2009).

20. Kim, Y.-J. et al. PTEN Modulates miR-21 Processing via RNA-Regulatory Protein RNH1. PLoS ONE 6, (2011).

21. Chennupati, V. et al. Ribonuclease inhibitor 1 regulates erythropoiesis by controlling GATA1 translation. J. Clin. Invest. 128, 1597–1614 (2018).

22. Hedberg-Oldfors, C. et al. Ribonuclease inhibitor 1 (RNH1) deficiency cause congenital cataracts and global developmental delay with infection-induced psychomotor regression and anemia. Eur J Hum Genet 1–8 (2023) doi:10.1038/s41431-023-01327-7.

23. Sarangdhar, M. A., Andina, N. & Allam, R. Human molecular genetics sheds light on the physiological significance of ribonuclease inhibitor (RNH1). Eur J Hum Genet 1–3 (2023) doi:10.1038/s41431-023-01362-4.

24. Choesmel, V. et al. Impaired ribosome biogenesis in Diamond-Blackfan anemia. Blood 109, 1275–1283 (2006).

25. Flygare, J. et al. Human RPS19, the gene mutated in Diamond-Blackfan anemia, encodes a ribosomal protein required for the maturation of 40S ribosomal subunits. Blood 109, 980–986 (2006).

26. Singh, S. A. et al. p53-Independent Cell Cycle and Erythroid Differentiation Defects in Murine Embryonic Stem Cells Haploinsufficient for Diamond Blackfan Anemia-Proteins: RPS19 versus RPL5. PLoS ONE 9, e89098 (2014).

27. Robledo, S. et al. The role of human ribosomal proteins in the maturation of rRNA and ribosome production. RNA 14, 1918–1929 (2008).

28. Liu, J., Xu, Y., Stoleru, D. & Salic, A. Imaging protein synthesis in cells and tissues with an alkyne analog of puromycin. Proc National Acad Sci 109, 413–418 (2011).

29. Guimaraes, J. C. & Zavolan, M. Patterns of ribosomal protein expression specify normal and malignant human cells. Genome Biol. 17, 236 (2016).

30. Elkordy, A. et al. Stress-induced tRNA cleavage and tiRNA generation in rat neuronal PC12 cells. J. Neurochem. 146, 560–569 (2018).

31. Oertlin, C. et al. Generally applicable transcriptome-wide analysis of translation using anota2seq. Nucleic Acids Res. 47, gkz223- (2019).

32. Thoreen, C. C. et al. A unifying model for mTORC1-mediated regulation of mRNA translation. Nature 486, 109–113 (2012).

33. Harding, H. P. et al. An Integrated Stress Response Regulates Amino Acid Metabolism and Resistance to Oxidative Stress. Mol. Cell 11, 619–633 (2003).

34. Quesne, J. P. C. L., Spriggs, K. A., Bushell, M. & Willis, A. E. Dysregulation of protein synthesis and disease. J. Pathol. 220, 140–151 (2010).

35. Lin, X. et al. Enhancer-Driven lncRNA BDNF-AS Induces Endocrine Resistance and Malignant Progression of Breast Cancer through the RNH1/TRIM21/mTOR Cascade. Cell Rep 31, 107753 (2020).

36. Kahvejian, A., Svitkin, Y. V., Sukarieh, R., M’Boutchou, M.-N. & Sonenberg, N. Mammalian poly(A)-binding protein is a eukaryotic translation initiation factor, which acts via multiple mechanisms. Gene Dev 19, 104–113 (2005).

37. Mangus, D. A., Evans, M. C. & Jacobson, A. Poly(A)-binding proteins: multifunctional scaffolds for the post-transcriptional control of gene expression. Genome Biol 4, 223–223 (2003).

38. Wells, S. E., Hillner, P. E., Vale, R. D. & Sachs, A. B. Circularization of mRNA by Eukaryotic Translation Initiation Factors. Mol. Cell 2, 135–140 (1998).

39. Cockman, E., Anderson, P. & Ivanov, P. TOP mRNPs: Molecular Mechanisms and Principles of Regulation. Biomolecules 10, 969 (2020).

40. Tcherkezian, J. et al. Proteomic analysis of cap-dependent translation identifies LARP1 as a key regulator of 5’TOP mRNA translation. Genes Dev. 28, 357–371 (2014).

41. Li, L. et al. Ribonuclease inhibitor up-regulation inhibits the growth and induces apoptosis in murine melanoma cells through repression of angiogenin and ILK/PI3K/AKT signaling pathway. Biochimie 1–12 (2014) doi:10.1016/j.biochi.2014.04.007.

42. Li, S. & Hu, G. Emerging role of angiogenin in stress response and cell survival under adverse conditions. J. Cell. Physiol. 227, 2822–2826 (2012).

43. Ivanov, P., Emara, M. M., Villen, J., Gygi, S. P. & Anderson, P. Angiogenin-induced tRNA fragments inhibit translation initiation. Mol. Cell 43, 613–623 (2011).

44. Moroianu, J. & Riordan, J. F. Nuclear translocation of angiogenin in proliferating endothelial cells is essential to its angiogenic activity. Proc. Natl. Acad. Sci. U.S.A. 91, 1677–1681 (1994).

45. Andina, N. D. et al. Higher Vertebrate Specific Gene Ribonuclease Inhibitor (RNH1) Is Essential for Adult Hematopoietic Stem Cell Function and Cell Cycle Regulation. Blood 134, 273–273 (2019).

46. Bianconi, E. et al. An estimation of the number of cells in the human body. Ann. Hum. Biol. 40, 463–471 (2013).

47. Goo, S. M. & Cho, S. The Expansion and Functional Diversification of the Mammalian Ribonuclease A Superfamily Epitomizes the Efficiency of Multigene Families at Generating Biological Novelty. Genome Biol. Evol. 5, 2124–2140 (2013).

48. Bosco, D. A. Translation dysregulation in neurodegenerative disorders. Proc. Natl. Acad. Sci. 115, 12842–12844 (2018).

49. Ruggero, D. Translational Control in Cancer Etiology. Cold Spring Harb. Perspect. Biol. 5, a012336 (2013).

50. Holler, N. et al. Two Adjacent Trimeric Fas Ligands Are Required for Fas Signaling and Formation of a Death-Inducing Signaling Complex. *Mol*. Cell. Biology 23, 1428–1440 (2003).

51. Kolly, C., Zakher, A., Strauss, C., Suter, M. M. & Müller, E. J. Keratinocyte transcriptional regulation of the human c-Myc promoter occurs via a novel Lef/Tcf binding element distinct from neoplastic cells. FEBS Lett. 581, 1969–1976 (2007).

52. Papin, S. et al. The SPRY domain of Pyrin, mutated in familial Mediterranean fever patients, interacts with inflammasome components and inhibits proIL-1beta processing. Cell Death Differ. 14, 1457–1466 (2007).

53. Sanchez-Taltavull, D., et al. Bayesian correlation is a robust gene similarity measure for single-cell RNA-seq data. NAR Genom. Bioinform. 2, lqaa002 (2020).

54. Goncalves, K. A. et al. Angiogenin Promotes Hematopoietic Regeneration by Dichotomously Regulating Quiescence of Stem and Progenitor Cells. Cell 166, 894–906 (2016).

55. Wang, L., Wang, S. & Li, W. RSeQC: quality control of RNA-seq experiments. Bioinformatics 28, 2184–2185 (2012).

56. Okonechnikov, K., Conesa, A. & García-Alcalde, F. Qualimap 2: advanced multi-sample quality control for high-throughput sequencing data. Bioinformatics 32, 292–294 (2016).

57. Kim, D., Langmead, B. & Salzberg, S. L. HISAT: a fast spliced aligner with low memory requirements. Nat. Methods 12, 357–360 (2015).

58. Liao, Y., Smyth, G. K. & Shi, W. featureCounts: an efficient general purpose program for assigning sequence reads to genomic features. Bioinformatics 30, 923–930 (2014).

59. Love, M. I., Huber, W. & Anders, S. Moderated estimation of fold change and dispersion for RNA-seq data with DESeq2. Genome Biol. 15, 31 (2014).

60. Alexa, A., Improv, J. R. B. & 2009. Gene set enrichment analysis with topGO. bioconductor.riken.jp.

61. Tyanova, S., Temu, T. & Cox, J. The MaxQuant computational platform for mass spectrometry-based shotgun proteomics. Nat Protoc 11, 2301–2319 (2016).

62. Katoh, K. & Standley, D. M. MAFFT multiple sequence alignment software version 7: improvements in performance and usability. Mol. Biol. Evol. 30, 772–780 (2013).

