## Supplementary Figures for "Ribonuclease Inhibitor and Angiogenin collaboratively regulate cell-type-specific global translation"

### Extended Data Fig. 1

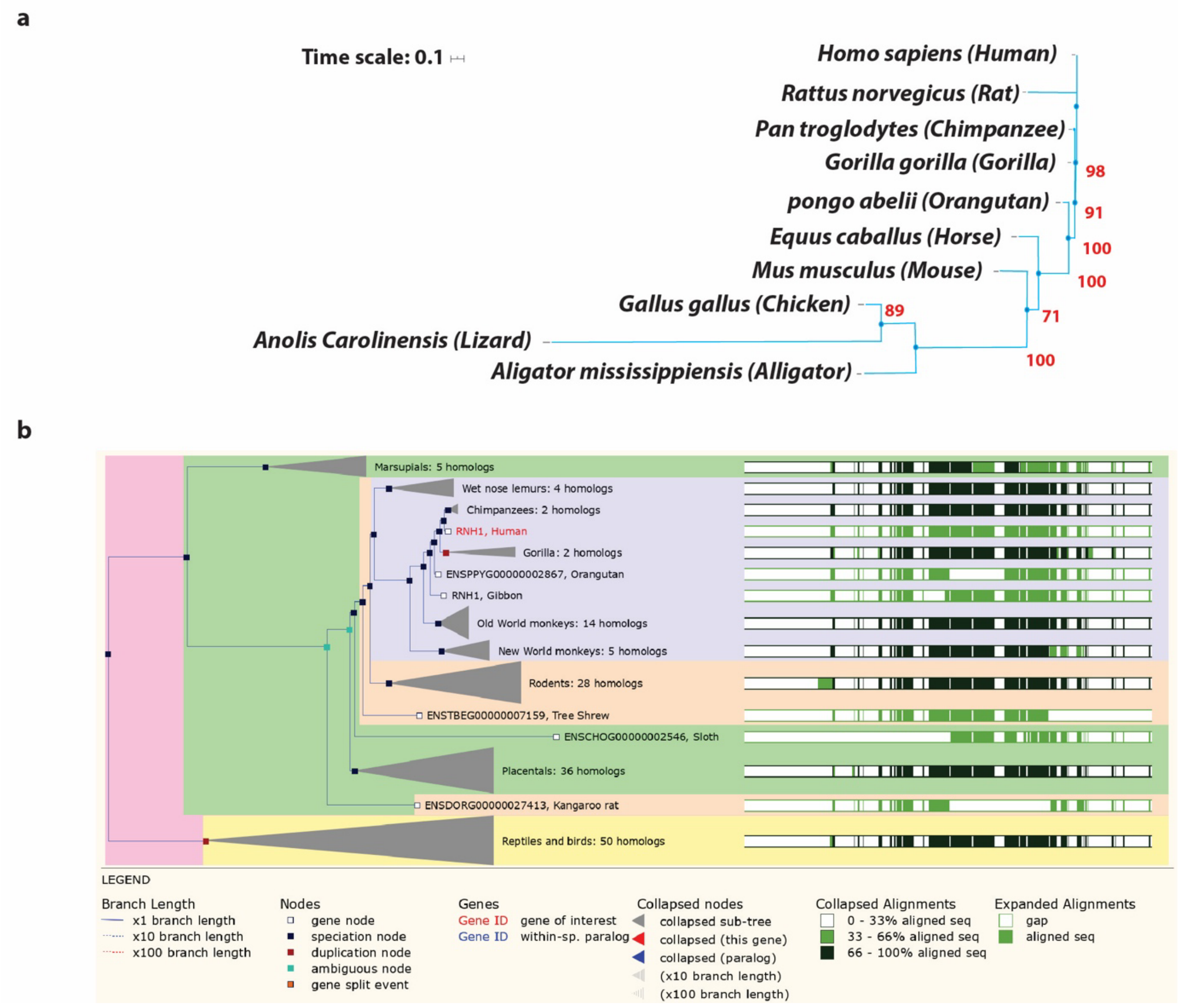


**Figure S1. *RNH1* Gene Evolved Only in Higher Vertebrates**

**(a)** Phylogenetic tree representing the evolutionary relationships between vertebrate RNH1 suggesting that it has diverged minimally among different subclasses of mammals. RNH1 DNA sequence alignments were made using MAFFT. A maximum likelihood phylogenetic tree was generated using IQ-Tree with 1000 bootstrap replicates. Bootstrap values >70% are shown in red. **(b)** Ensembl gene tree for RNH1 generated by the Gene Orthology/Paralogy prediction method. Gene trees are constructed using one representative protein for every gene in every species in Ensembl. The figure shows the maximum likelihood phylogenetic tree representing the evolutionary history of *RNH1* gene.

**Extended Data Fig. 2**

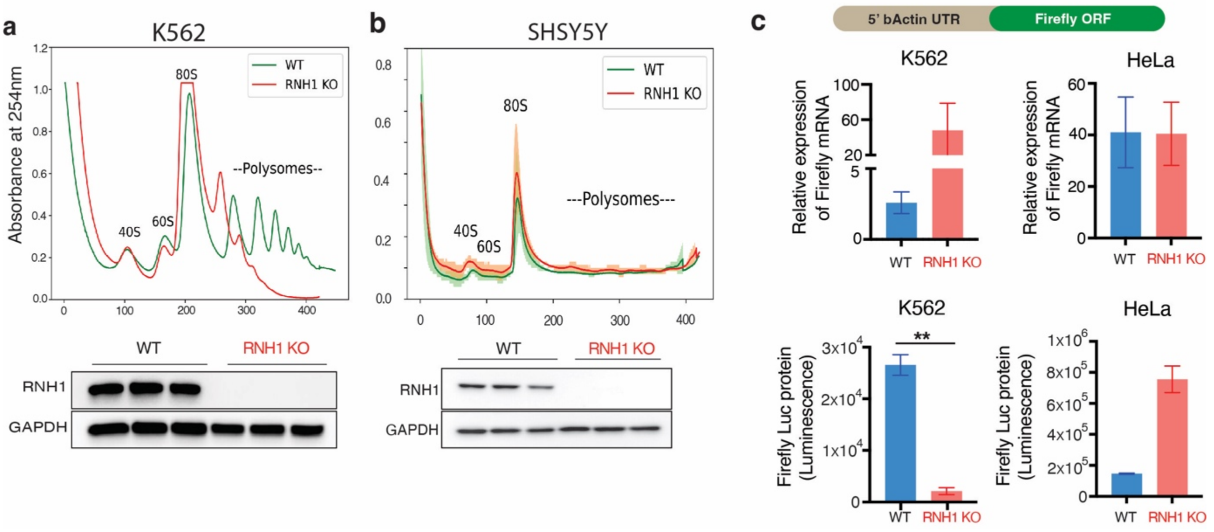


**Figure S2. Loss of RNH1 decreases translation in hematopoietic cells but not in non-hematopoietic cells**

(**a and b**) Sucrose gradient polysome profiles for WT and corresponding RNH1 KO of K562 or SHSY5Y cells (N=3). Arrow shows the direction of the sucrose gradient from low to high density. Mean value of absorbance from three independent experiments plotted with the standard deviation (upper panel). Total protein lysates of WT and RNH1 KO cells were analyzed by western blot with the indicated antibodies. Blots are representative of three independent experiments (lower panel). (**c**) Schematics of luciferase expressing plasmid with beta actin (ACTIN B) 5`UTR (upper panel). WT or RNH1 KO of HeLa or K562 cells were transfected with luciferase expressing plasmid. Cells were analysed for firefly mRNAs by qRT-PCR, normalized to 18S rRNA expression and luciferase protein expression by luciferase assay. Data are shown as mean ± SEM and are representative of 3 independent experiments.

**Extended Data Fig. 3**

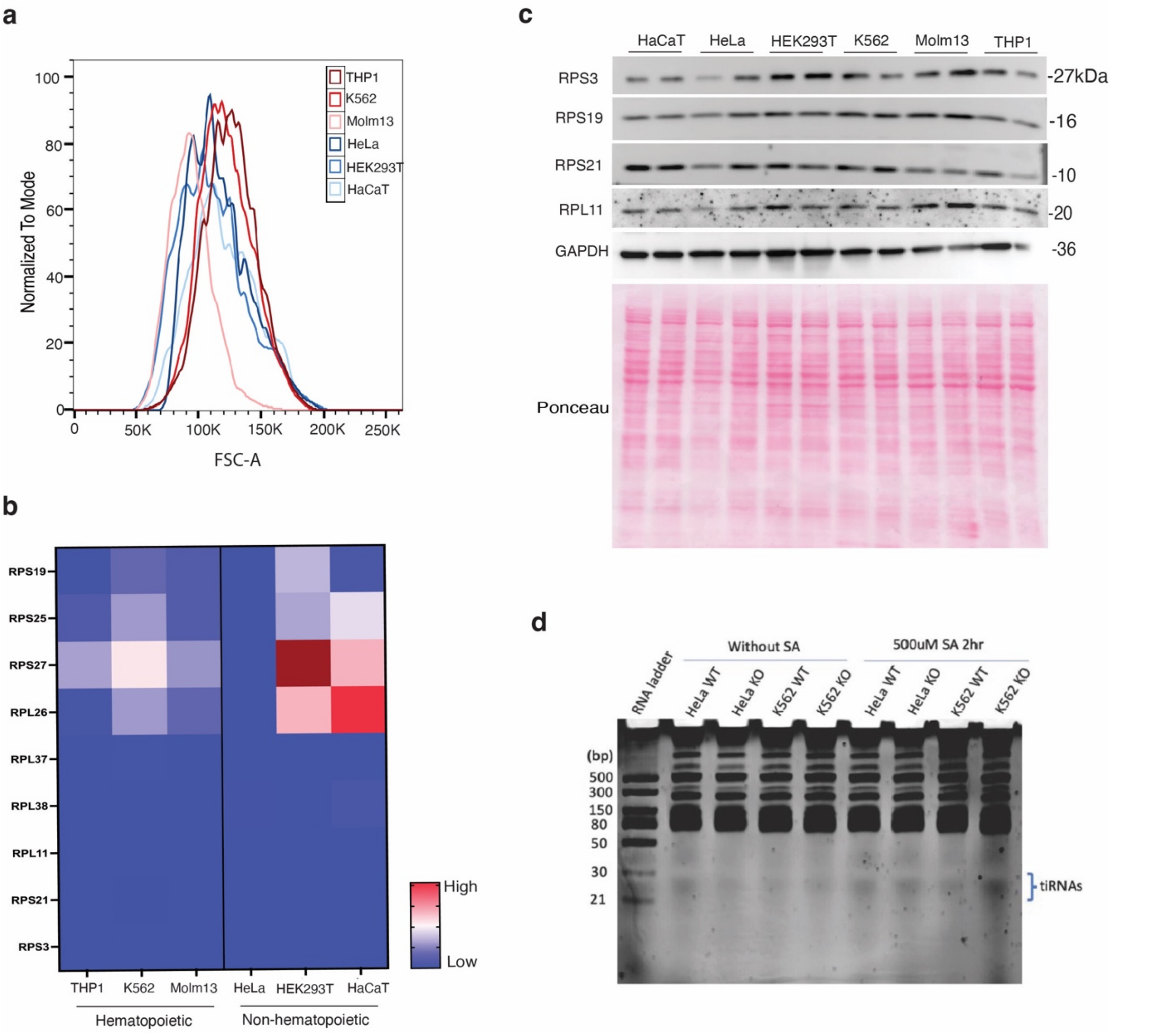


**Figure S3. RNH1 mediated translation specificity is not dependent on cell size and RPs gene expression**

(**a**) flow cytometry analysis for forward scatter (FSC) to evaluate cell size from different hematopoietic and non-hematopoietic cells. (**b**) Heatmap showing qRT-PCR results of mRNA relative expression of different ribosomal protein genes from hematopoietic and non-hematopoietic cell lines. mRNA levels were normalized to beta-Actin mRNA. Results are representative of two independent experiments. (**c**) Total protein lysates of different hematopoietic and non-hematopoietic cell lines were analyzed by western blot with the indicated antibodies. Blots are representative of three independent experiments. (**d**) HeLa and K562 cells of WT or RNH1 KO conditions were collected after 2hr with or without stimulation by 500uM sodium arsenite (SA) and RNA was isolated. 3ug of RNA was mixed with RNA loading dye and heated for 5min, 60°C and loaded on Urea-PAGE for tiRNA separation. Gels are representative of 3 independent experiments.

**Extended Data Fig. 4**

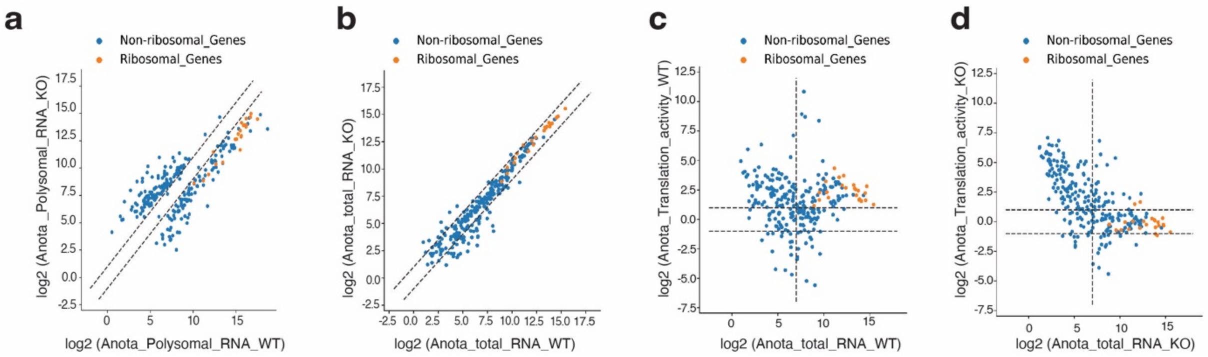


**Figure S4. RNH1 mediates RPs gene translation**

(**a and b**) Polysomal RNA-seq and total RNA-seq data of WT and RNH1 KO K562 cells were subjected to ANOTA analysis. The expression of ANOTA filtered ribosomal and non-ribosomal genes (Total 262 genes) with cutoff of padj. <0.1 in polysomal RNA-seq and total RNA-seq were selected and plotted. Dotted lines separate genes with log2(fold change) <1 and >1. (**c and d**) Plot of Log2 fold change of total RNA verses translational activity (TA) of ribosomal and non-ribosomal genes in WT (C) or RNH1 KO (D) samples. Horizontal dotted lines separate genes with log2(fold change) <1 and >1. Vertical line separate genes with arbitrary cutoff of >7< for log2(expression in total RNA-seq).

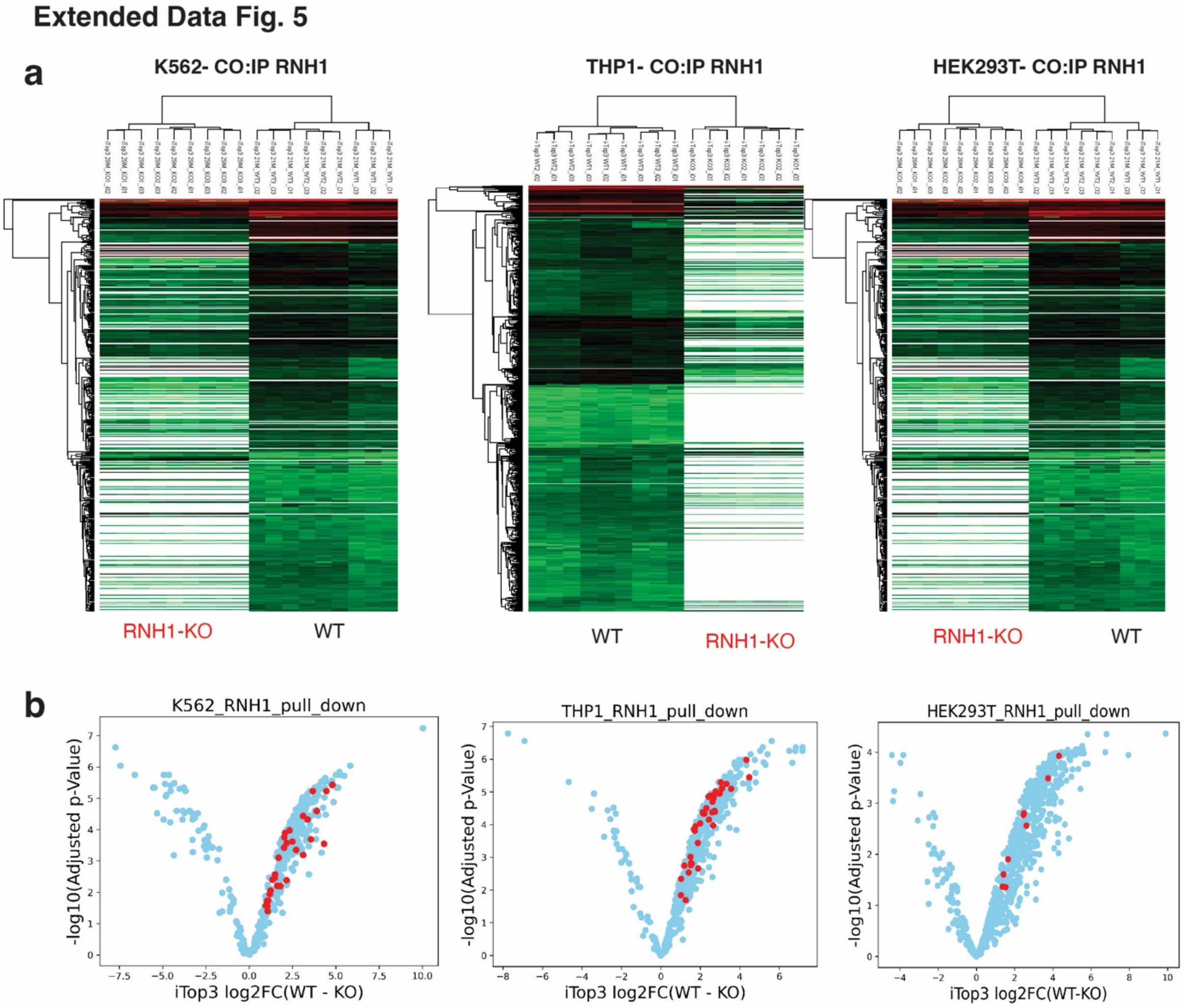


**Figure S5. RNH1 binds to Ribosomal proteins**

(**a**) Heatmap showing Cluster analysis with iTop3 values of RNH1 binding total proteome from WT THP1 or K562 or HEK29T cells compared with RNH1 KO corresponding cells. (probability of false positive < 0.1). (**b**) Volcano plot showing ribosomal proteins in red.

**Extended Data Fig. 6**

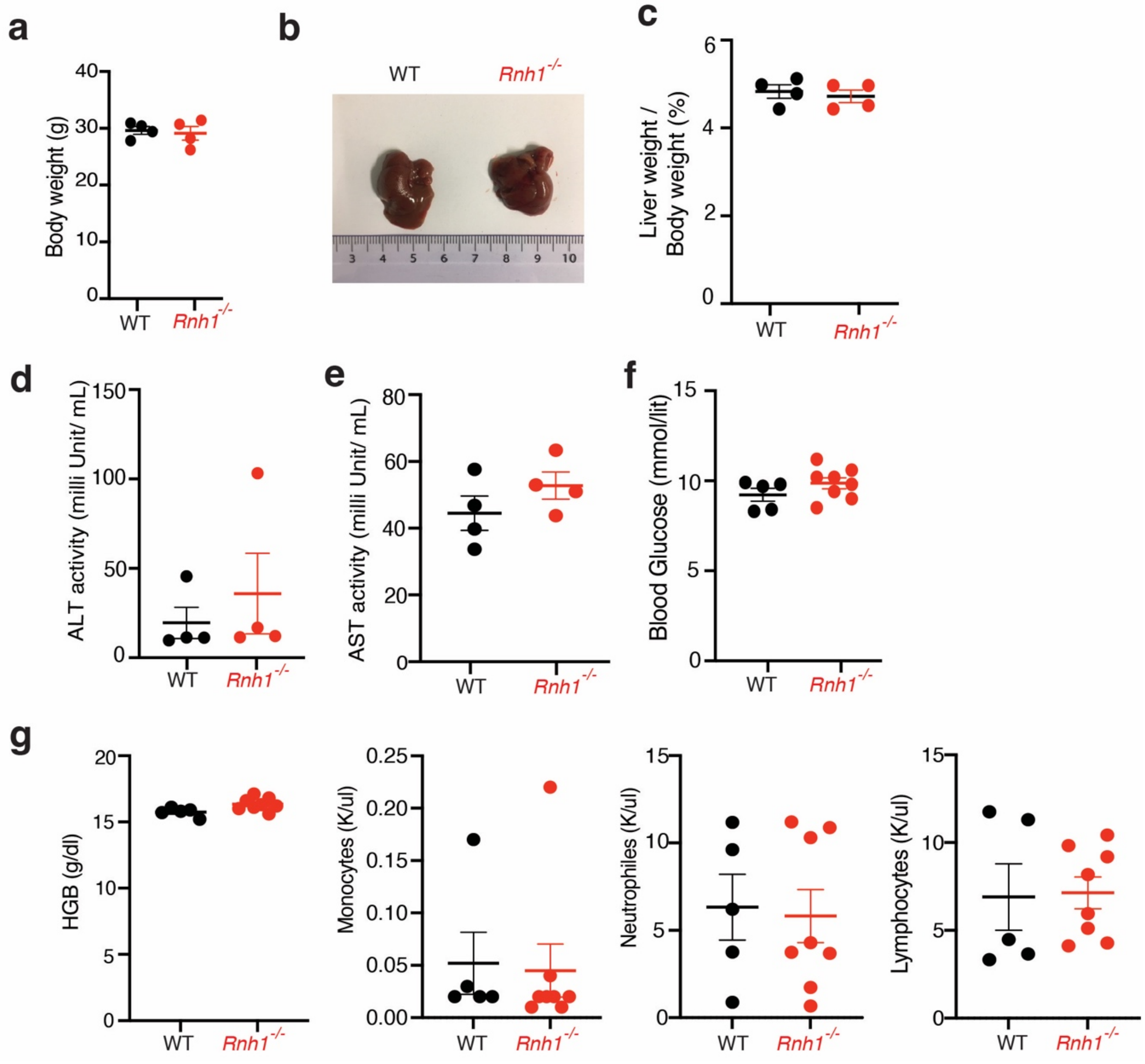


**Figure S6. Loss of RNH1 in liver doesn't affect the liver function and overall survival**

**(a)** Body weight of WT (*RNH1* $^{fl/fl}$) and *Rnh1*$^{-/-}$ (*RNH1* $^{fl/fl}$*, Alb-Cre*$^{+}$) mice at 14 weeks of age (N=4). **(b and c)** Representative image of liver and liver weight of WT and *Rnh1*$^{-/-}$ mice at 14 weeks (n = 4 mice). (**d and e**) Serum ALT and AST levels of WT and *Rnh1*$^{-/-}$ mice at 14 weeks (n = 4 mice). **(f and g)** Blood glucose levels and peripheral blood (PB) counts of neutrophils, granulocytes, lymphocytes, and haemoglobin (HBG) concentration in WT and *Rnh1*$^{-/-}$ mice at 14 weeks (n = 5 - 8 mice). All the data are shown as mean ± SEM.
